# Generating and navigating single cell dynamics via a geodesic bridge between nonlinear transcriptional and linear latent manifolds

**DOI:** 10.64898/2026.03.31.715478

**Authors:** Junchao Zhu, Zhenyi Zhang, Yuhao Sun, Hao Dai, Han Wen, Peijie Zhou, Luonan Chen

## Abstract

Time-series single-cell RNA sequencing (scRNA-seq) captures cellular processes as sparse and unpaired snapshots, limiting our ability not only to reconstruct continuous cell state transitions, but also to *navigate* between states in a controlled and interpretable manner. Here we present **GeoBridge**, a framework modeling cellular dynamics as geodesic trajectories on the transcriptional manifold based on isometric geodesic theory. By learning the geodesic bridge mapping, the method theoretically and computationally transforms time-varying nonlinear transcriptional geodesics (original nonlinear manifold) into constant-velocity straight-line geodesics (latent linear manifold). In the learned geodesic space, continuous interpolation becomes biologically meaningful, enabling reconstruction of unobserved intermediate states and efficient navigation between distinct cellular phenotypes at a single-cell resolution. By mapping interpolated trajectories back to the original gene expression space, GeoBridge recovers smooth transcriptional programs that are robust to noise and snapshot sparsity. Leveraging the derived geodesic potentials, GeoBridge further infers pseudo-temporal trajectories with superior fidelity compared to mainstream approaches from single-snapshot scRNA-seq data without temporal annotation, and directly identifies genes that drive progression along transition paths. We demonstrate the utility of GeoBridge across diverse systems, where GeoBridge resolves EMT–MET progression in cancer stem cells and identifies stage-specific modules as well as the branching cell-fate dynamics in human pluripotent stem cells with higher reconstruction accuracy than state-of-art methods. More importantly, GeoBridge supports single-cell fate navigation in multi-target hematopoietic lineages, allowing neutrophil-biased cells to be virtually guided toward mast-cell fates along biologically plausible paths. Together, GeoBridge establishes a principled method that transforms sparse single-cell measurements into a continuous, controllable landscape for the reconstruction, navigation and manipulation of cellular state transitions.

## Introduction

Understanding how cell states evolve continuously during differentiation or disease progression is central to controlling and engineering biological outcomes . Recent advances in single-cell RNA sequencing (scRNA-seq) have provided the foundation for comprehensive profiling of cellular states heterogeneity^1^. However, for continuously evolving biological processes, such as stem cell differentiation or phenotypic plasticity, scRNA-seq can only capture discrete snapshots at a limited number of time points. Furthermore, because scRNA-seq is terminal and destructive, cells at different time points cannot be paired directly. As a result, our understanding of these processes lacks a continuous and holistic temporal view, making it difficult to extract underlying biological dynamic principles and challenging to infer smooth, biologically meaningful and navigable trajectories between cell states^2–6^.

To address these challenges, several pioneering approaches have been developed to approximate continuous cellular transitions from static data. Pseudotime methods^7–9^ infer an ordering of cells, providing insight into branching structures. More recently, RNA velocity^10^ analysis has extended this idea by estimating short-term future transcriptional states based on the relative abundance of unspliced and spliced mRNAs, thereby introducing a pseudo-dynamic layer on top of static transcriptomes. Furthermore, approaches based on Dynamic Optimal Transport (DOT)^11–13^ have been widely adopted to infer cell-to-cell correspondences and continuous trajectories under the principle of minimizing total transport energy, provided an action-minimization framework for modeling developmental pathways. Existing methods such as TrajectoryNet^14^, TIGON^15^, and DeepRUOT^16^ typically compute the cellular action under the assumption of an Euclidean metric, meaning each gene contributes equally to the distance between two cells. However, under the manifold hypothesis, high-dimensional biological data often reside near a low-dimensional differentiable manifold^17^, the “distance” between cells becomes a nonlinear function of gene expression, reflecting that different genes contribute unequally depending on biological context. In this context, the minimal-action path between two points should correspond to a geodesic under the associated Riemannian metric of the manifold^18–20^, with the velocity norm computed according to this metric. Assuming an Euclidean metric in this curved, intrinsic cell-state space leads to geometric distortion of data, resulting in trajectories that are *non-geodesic* and thus inaccurate estimations of developmental speed, fate probability, and distance between states, which clearly contradicts the biological reality, e.g. lineage-specific genes are far more informative than housekeeping genes for defining differentiation paths.

However, in high-dimensional and noisy single-cell data, directly computing exact geodesic distances in the original space is practically infeasible. To circumvent this, recent works such as MIOFlow^21^ employ diffusion distance in the original gene-expression manifold as a computationally tractable surrogate for geodesic distance. While effective in practice, such approximations blur the connection between inferred trajectories and the underlying biological geometry, limiting interpretability and control of the single-cell dynamics.

To overcome these limitations, we introduce GeoBridge, a data-driven framework that constructs a bijective geodesic bridge between the native nonlinear transcriptional manifold and a latent flat Euclidean manifold. GeoBridge’s core biological motivation is to preserve the intrinsic “biological distance” between cells by modeling cellular dynamics as geodesic trajectories on the transcriptional manifold based on our isometric geodesic theory. GeoBridge employs an Invertible Neural Network (INN) ^22^ to learn an isometric mapping, which is mathematically proven to preserve intrinsic geodesic distances (Figure 1 and Supplmentary Note 1). Rather than explicitly computing geodesics in the original space, the method exploits the property of flat manifolds ^23–28^: geodesics correspond to constant-velocity straight lines. By enforcing a constant-velocity linear constraint in the latent space, GeoBridge implicitly drives the learned metric toward an ideal Euclidean geometry, without requiring direct access to geodesic distances in the original manifold.

**Fig 1.**
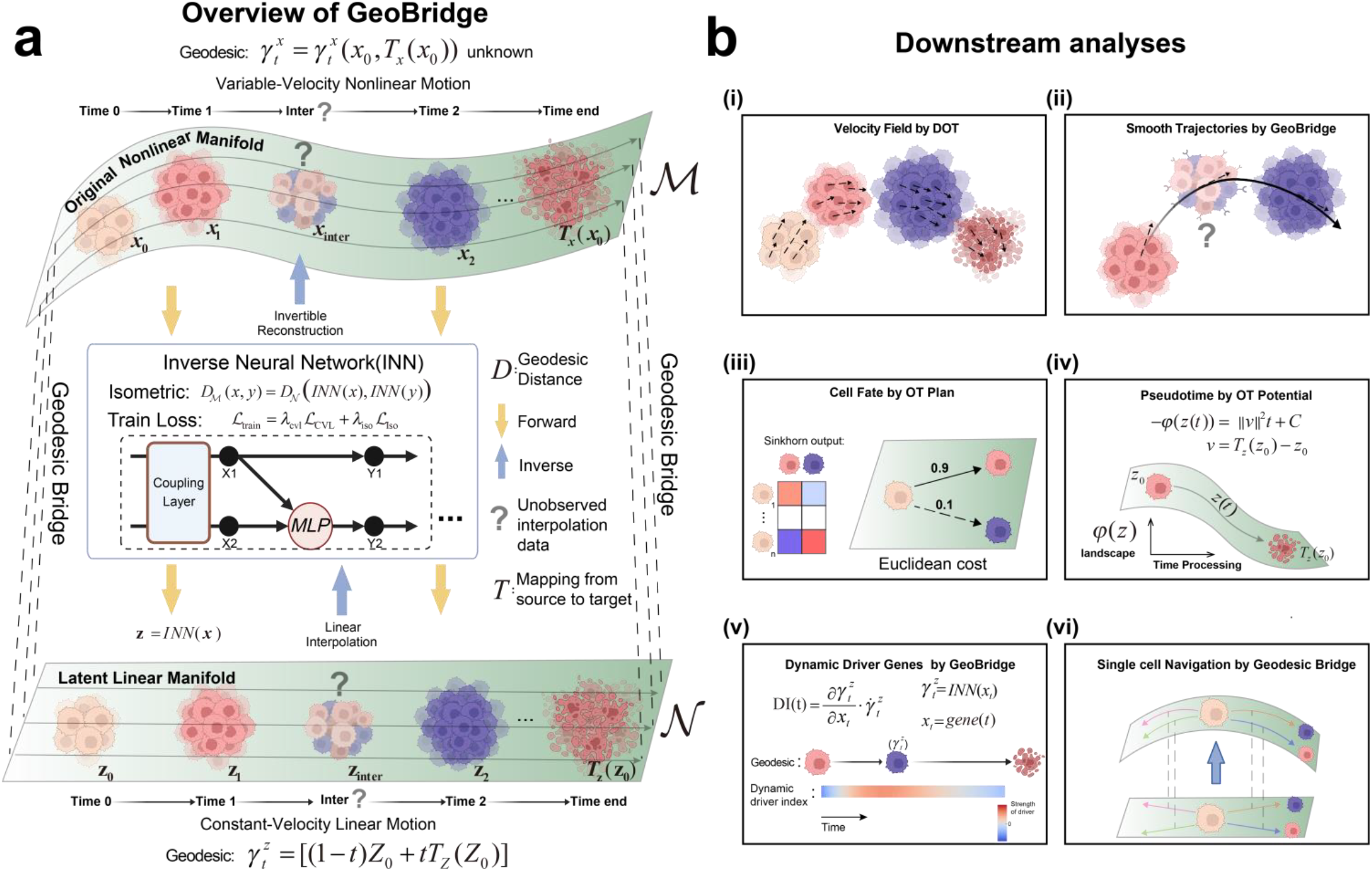
Illustrative overview of GeoBridge for Time-series single-cell data. **a**, Time-series single-cell data are assumed to lie on a Riemannian (original nonlinear) manifold M, where the true biological progression follows a minimal-action geodesic path. GeoBridge employs an invertible neural network (INN) to map this gene-expression manifold *M* onto a flat forward Euclidean (embedded linear) manifold *N* through a constant-velocity linear loss (ℒCVL). Constant-velocity straight-line geodesics in *N* then were ensured one-to-one correspondence to original nonlinear geodesics in *M* through local isometry regularization (ℒ_Iso_), allowing accurate reconstruction of continuous biological evolution via linear interpolation and inverse mapping. **b**, Downstream analyses derived from the latent linear manifold learned by GeoBridge. The computational details are given in Downstream analyses in Methods. (i–ii) Inverse-mapped linear geodesics yield single-cell velocity fields and progression trajectories. (iii) Single-cell fate is determined by optimal transport weights. The cost function is Euclidean geodesic distance in N. (iv) Kantorovich potentials computed provide pseudotime estimates linear with respect to real time parameter. (v) Single-gene-level dynamic driver index (DI) is derived as dot products between trajectory expression gradients and velocities. (vi) Given a starting cell and any target cell, GeoBridge generates the geodesic navigation path by constructing and inverse-mapping straight-line geodesic.

The established geodesic bridge simplifies complex, nonlinear cell state transitions into straight-line paths in latent space. Intermediate states can therefore be generated analytically through closed-form linear interpolation. Mapping these interpolations back through the inverse transformation reconstructs geodesic trajectories in gene expression space, corresponding to the theoretically shortest, minimum-action paths between cellular states. In this way, GeoBridge enables biologically meaningful interpolation, trajectory design, and state navigation at single-cell resolution.

Taken together, the proposed bijective geodesic bridge eliminates the need for intractable geometric computations on the original single-cell gene expression manifold, providing a simulation-free generative model^29–31^ of cellular dynamics rooted in precise geometry. Unlike relevant methods (e.g., TrajectoryNet, TIGON, DeepRUOT, MIOFlow) which rely on computationally intensive numerical integration via Neural ODEs and often require preliminary dimensionality reduction, GeoBridge performs interpolation via an explicit, closed-form linear combination and can operate directly on high-dimensional inputs. By establishing a latent linear manifold with preserved geometric properties, we apply GeoBridge to multiple real scRNA-seq datasets to demonstrate its capability for 1) precise cellular dynamic characterization, 2) accurate single-cell fate prediction, 3) driver gene identification, as well as 4) controllable geodesic navigation between arbitrary cellular states.

## Results

### Overview of GeoBridge and isometric geodesic theory

We suppose that time-series single-cell transcriptomic data are distributed over a Riemannian manifold *M* embedded in a high-dimensional space^17^. The true biological progression trajectory corresponds to the path of minimal action on *M*; according to dynamic optimal transport (DOT) theory, this path is a geodesic, i.e., the shortest path in the sense of the Riemannian metric. Biologically, this geodesic represents the most feasible transition path between cell states, minimizing the number of transcriptional changes and aligning with the gradual reprogramming of gene regulatory networks observed in development. In the original data (nonlinear) manifold, however, the Riemannian metric of *M* is unknown, and directly generating geodesics via data-driven methods often incurs substantial computational cost and reduced accuracy.

To address this, we employ an Invertible Neural Network (INN) to map the data from *M* to a flat (linear) manifold *N* endowed with an Euclidean metric (Fig.1a and Methods) based on our isometric geodesic bridge theory, i.e. bijective diffeomorphism and geodesic bridge theorem (Theorem 2 in SI). On *N*, the minimal action path is a constant-velocity linear trajectory. This enables us to reconstruct the biological progression trajectories in the original manifold by just performing linear interpolation at intermediate time points in *N* and inverse-mapping the interpolants to *M*.

Conceptually, our approach “straightens” complex nonlinear manifold geodesics into lines. Crucially, geodesics on *M* and *N* must be in one-to-one correspondence; to ensure this property, we introduce a local isometry regularization constraint (Equation (7)).

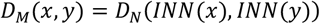

where *D*_*M*_ denotes the Riemann geodesic distance on the original (nonlinear) manifold *M, D*_*N*_ denotes the Euclidian geodesic distance on the forward mapped (embedded linear) manifold *N*, and *x* and *y* are neighboring cells on the *M. INN*(*x*) means to map the data *x* from *M* to *N*.

We further prove that, under this constraint, a constant-velocity straight-line geodesic in N, when mapped back via the inverse INN transformation, remains a geodesic in *M* (Theorem 2 in SI). From an intuitive perspective, preserving the local distance relationships ensures that the local geometric structure is undistorted, meaning that the relative expression patterns of biologically relevant genes are maintained so that the reconstructed trajectories reflect true transcriptional dynamics rather than artificial distortions. Under this condition, the shortest paths with respect to each manifold’s metric (geodesics) are maintained in a one-to-one correspondence.

Our ablation studies (Supplementary Fig. S1-S2) demonstrate that while a constant-velocity linear loss alone is sufficient to create a flat latent manifold, but without the isometric constraint, when straight-line paths from the latent manifold are mapped back to the original gene expression manifold, the generated intermediate cellular states can deviate substantially from the true data manifold, leading to biologically implausible states.

Based on this principle, inverse-mapped straight-line geodesics from *N* provide both the single-cell trajectories and the velocity fields in the original space (Fig.1b i–ii). Such a project is called a geodesic bridge, and our method is also named GeoBridge in this work. In static optimal transport problems, the key determinant is the choice of an appropriate cost function rather than the optimization method itself. Within the dynamic optimal transport framework, the cost between source and target domains is the geodesic length between them^32^. In our model, the geodesic length between two points in *M* is equivalent to the Euclidean distance between their mapped points in *N*. Thus, our method can be interpreted as learning the point–point metric or geodesic bridge in high-dimensional space. This formulation facilitates the solution of static OT problems, enabling us to estimate cell fate weights via the optimal transport plan (Fig.1b iii). Moreover, when the cost function is Euclidean distance, the Kantorovich potential (KP) of the optimal transport exhibits a linear relationship with the true temporal parameter (Equation (35)). We solve this relationship using the FISTA-OT method^33^ and employ the resulting KP as a pseudotime indicator for cells (Fig.1b iv).

Building upon GeoBridge, we additionally compute, at each time point, the gradient of the straight-line geodesic in *N* with respect to each gene’s expression in *M*, and take the dot product with the geodesic’s velocity components across directions. This yields the dynamic driver effect of each gene along the trajectory (Equation (40)), where a positive value indicates a driving influence on the trajectory, and a negative value indicates suppression (Fig.1b v).

Finally, we explore an extension: for any given cell, can we produce its geodesic trajectory to any target endpoint? We refer to this task as single-cell navigation (Fig.1b vi). For an arbitrary cell, we first construct the straight-line geodesic in *N* to the target cell, and then reconstruct its geodesic in *M* via inverse mapping through the INN.

### Reconstructing continuous cellular dynamics in EMT– MET progression

We applied GeoBridge to a time-series dataset capturing the developmental process of cancer stem cells (CSCs) across five time points (day 0–day 30)^14^. During this period, CSCs underwent epithelial–mesenchymal transition (EMT) and mesenchymal– epithelial transition (MET), ultimately differentiating into mesenchymal (Mes) or epithelial (Epi) states (Fig. 2a). To model these distinct lineage directions, we trained two independent INN models: the Mes-INN using data from day 0–day 18 and day 30 Mes cells, and the Epi-INN using day 0–day 18 data and day 30 Epi cells.

**Fig. 2.**
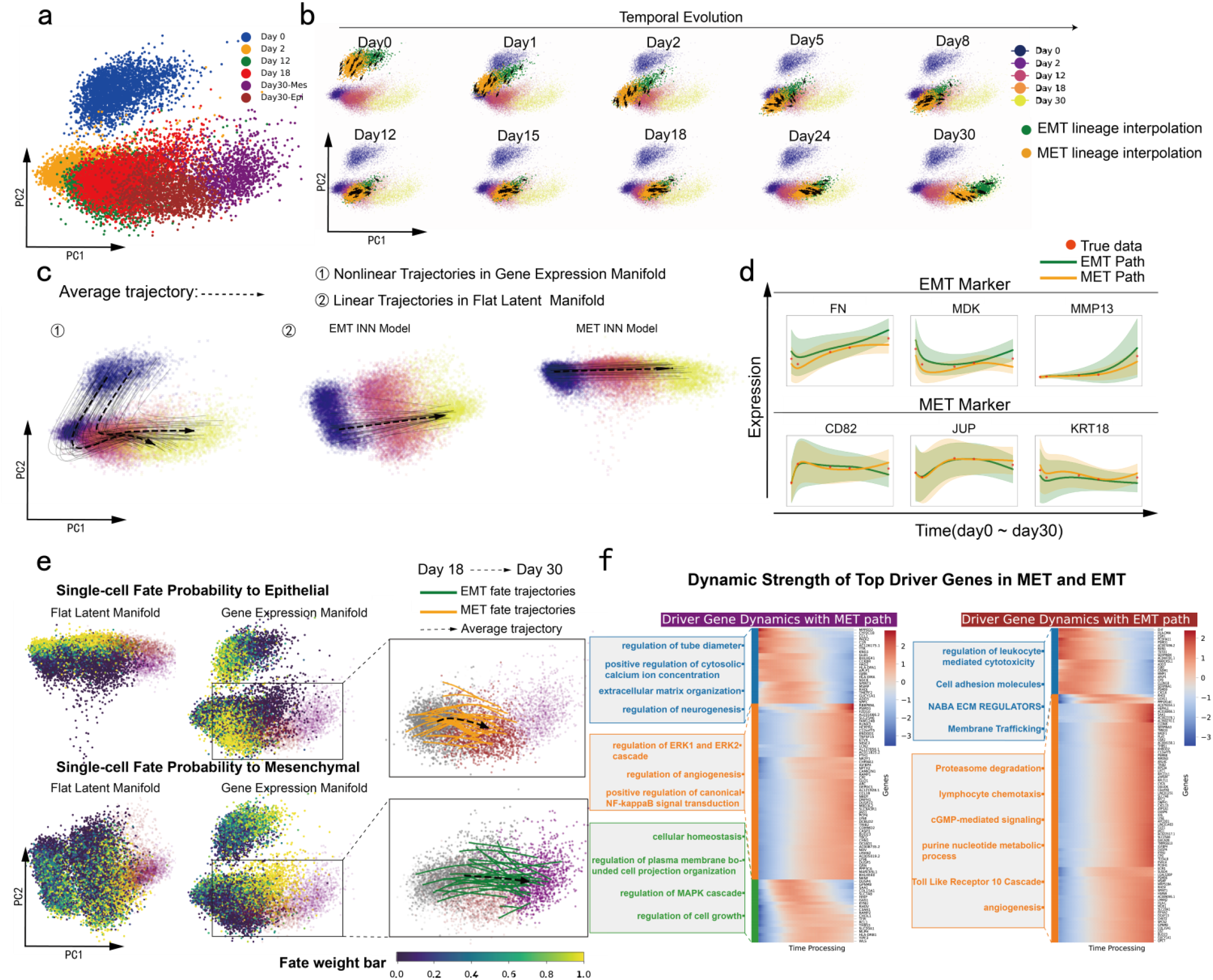
GeoBridge reconstructs cellular dynamics during EMT–MET progression. **a-c**,**e** The data is visualized in PCA embedding. **a**, Data of cancer stem cells (CSCs) showing in original gene-expression (nonlinear) manifold(day 0–30). Including EMT(Day30-Mes target) and MET(Day30-Epi target) branches. **b**, Reconstruction of intermediate cellular states in gene-expression manifold between day 0 and day 30. Including EMT lineage(green) and MET lineage(orange). The arrows indicate the velocities of ten randomly selected cells from each lineage. **c**, Thirty single-cell trajectories along EMT and MET lineages in the gene-expression manifold(①); Corresponding straight-line geodesics in the flat forward (embedded linear) manifold(②) **d**. Differential dynamics Marker-gene expression of continuous EMT(green)/MET(orange) progression (FN, MDK, MMP13; CD82, JUP, KRT18). **e**, left. Epithelial and Mesenchymal fate-probability estimations for day 0–18 CSCs cells in the two manifolds. right. The 30 randomly selected trajectories from day 18 CSCs(grey) to day 30 Epithelial(orange) and Mesenchymal(green). **f**, Heatmap of Top 100 dynamic driver index highlight temporally distinct modules of regulatory genes along EMT and MET lineages. GO enrichment of driver genes showing early(blue), mid(green) and late(orange) programs in MET and early(blue), late(orange) programs in EMT.

Within each flat forward manifold, we computed the cell fate probabilities of day 0–day 18 cells toward Mes or Epi endpoints, thereby assigning individual cell fates (Supplementary Fig. S3 a– d, Methods). Linear interpolation between the initial (day 0 fate) and terminal (day 30) states in the latent linear manifold(Supplementary Fig. S9.a), followed by inverse mapping to the original gene expression manifold and PCA visualization, reconstructed unobserved middle data for both lineages (Fig. 2b).

To illustrate dynamic transitions, we traced 30 single-cell trajectories along each lineage and overlaid their mean paths (Fig. 2c ①). Corresponding linear geodesics in the latent manifold (Fig. 2c ②) revealed that straight lines in latent space become nonlinear geodesics in gene expression manifold—indicating that the INN preserves biological continuity while capturing transcriptional nonlinearity. Obviously, the model input were the top 1,999 highly variable genes so that the geodesics reconstructed by inverse-mapping were continuous modeling of gene-expression dynamics, exemplified by the time-resolved expression of EMT markers (FN, MDK, MMP13) and MET markers (CD82, JUP, KRT18) (Fig. 2d). Notably, EMT marker genes exhibited higher expression along the EMT trajectory, whereas MET marker genes were upregulated along the MET trajectory, consistent with their established lineage identities. This concordance reveal that the inverse-mapped geodesics capture biologically meaningful transcriptional states.

Fate-probability mapping further distinguished Mes- and Epi-fated subpopulations within day 0–day 18 cells (Fig. 2e). Using day 18 CSCs as branch points, we traced trajectories linking Mes-fated cells to day 30 Mes cells and Epi-fated cells to day 30 Epi cells (Fig. 2e, right). To identify molecular determinants of lineage bias, we computed gene-specific dynamic driver indices that quantify contributions to EMT or MET progression from day 18 to day 30. Driver strength correlated strongly with gene expression (Supplementary Fig. S3e). The top 100 driver genes exhibited temporally distinct activity patterns: three temporal modules (early, mid, late) in MET and two (early, late) in EMT (Fig. 2f). GO enrichment analyses^34^ (Supplementary Fig. S3f–g) revealed that early MET drivers were enriched for extracellular matrix organization^35^, calcium signaling^36^, and regulation of tube diameter etc, indicating microenvironmental reorganization and polarity establishment; mid-stage drivers were enriched for MAPK^37^ regulation, cell growth and homeostasis etc, consistent with epithelial stabilization; late drivers mediated regulation of ERK cascades^38^ and NF-κB signal transduction^38^, and angiogenic remodeling^39^ etc, reflecting proliferative and metabolic adaptation. In contrast, early EMT drivers were enriched in leukocyte-mediated cytotoxic regulation^40^, adhesion^40,41^, and matrix remodeling^35^ etc, facilitating motility acquisition, whereas late drivers activated proteasome degradation, cGMP signaling, Toll-like receptor, and chemotaxis pathways, reflecting heightened metabolic and immune crosstalk supporting tissue remodeling and metastatic competence.

### Resolving hPSCs lineages toward SC-β and SC-EC

We further applied GeoBridge to an in vitro induction system modeling the differentiation of human pluripotent stem cells (hPSCs) into stem cell–derived β cells (SC β) and enterochromaffin like cells (SC EC)^42^. The dataset comprised eight time points (day 0–day 7) and six cell types (Fig. 3a). To capture the dynamic processes along the two lineage directions, we trained two independent invertible neural network (INN) models: β INN, using early stage stem cell data together with terminal SC β cells, and EC INN, using early stage stem cell data together with SC EC cells.

**Fig. 3.**
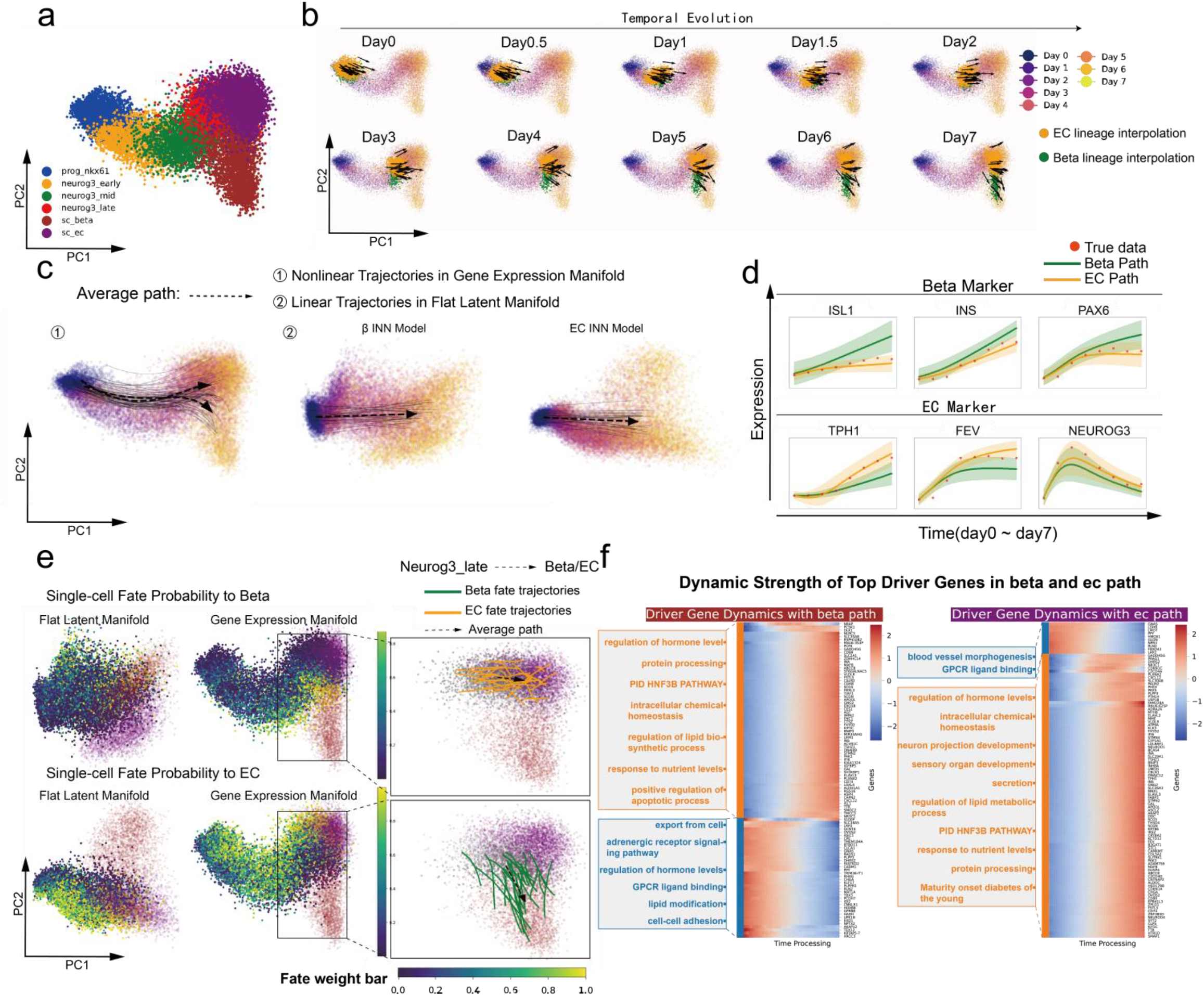
GeoBridge resolves cellular dynamics across pancreatic endocrine lineages. **a–c**,**e** The data are visualized in PCA embedding. **a**, Time-series Single-cell pancreatic endocrine lineage transcriptomic data showing the gene-expression manifold. HPSCs gradually differentiate toward β-cell (brown) and enterochromaffin cell (purple) fates. **b**, Reconstruction of intermediate cellular states in the gene-expression manifold between early Prog_NKX6.1 stage and mature β/EC cells. Including β-cell lineage (green) and EC lineage (orange). The arrows indicate the velocities of ten randomly selected cells from each lineage. **c**, Thirty single-cell trajectories along β and EC lineages in the gene-expression manifold (①); corresponding straight-line geodesics in the flat forward manifold (②) derived from β-INN and EC-INN models. **d**, Dynamics of marker-gene expression illustrating continuous β (green)/EC (orange) progression (ISL1, INS, PAX6; TPH1, FEV, NEUROD3). **e**, left, Fate-probability estimations for progenitor cells along β and EC manifolds. right, Thirty randomly selected trajectories connecting progenitors in NEUROG3_late state(grey) to terminal β (green) and EC (orange) fates. **f**, Heatmap of Top 100 dynamic driver indices highlighting temporally distinct regulatory modules along β and EC lineages. GO enrichment of driver genes showing early (blue), mid (green) and late (orange) programs in β differentiation, and early (blue), late (orange) programs in EC differentiation.

Within each flat forward manifold, we calculated the single cell fate probability toward β or EC endpoints to define their potential differentiation direction (Supplementary Fig. S4a–d, Methods). We then performed linear interpolation between the Prog_NKX6.1 stage (β/EC fate) and the terminal SC β or SC EC states within the forward manifold(Supplementary Fig. S9.b), followed by inverse mapping back to the original nonlinear manifold or gene expression manifold and PCA projection for visualization (Fig. 3b).

To reveal continuous dynamic changes, we randomly selected 30 single cell trajectories along β and EC pathways and plotted their mean trajectories (Fig. 3c①). Correspondingly, the straight geodesics and their mean paths in the forward manifold (Fig. 3c②). Using the top 2,000 highly variable genes as input, this process effectively modeled gene expression dynamics from discrete points to continuous transitions. We further illustrated time resolved expression of β lineage marker genes (INS, ISL1, NEUROG3) and EC lineage marker genes (TPH1, FEV, PAX6) across days 0–7 (Fig. 3d).

In both forward and gene-expression manifold, fate distributions clearly separated β biased and EC biased populations (Fig. 3e). Starting from the intermediate NEUROG3_late state, we traced single cell trajectories connecting β biased cells to SC β cells and EC biased cells to SC EC cells (Fig. 3e, right).

Finally, we quantified gene specific driver indices along SC β and SC EC differentiation trajectories to assess their promoting or inhibitory effects. Driver strength showed a strong correlation with expression level (Supplementary Fig. S4e). Among the top 100 genes ranked by mean driver index, β pathway and EC path drivers were classified into early (blue), and late (orange) phases (Fig. 3f). GO enrichment^34^ (Supplementary Fig. S4f–g) further revealed distinct regulatory programs. During differentiation toward SC β cells, early stage enrichment involved molecular transport, GPCR signaling, and hormone regulation^43,44^ etc, indicating the initiation of endocrine function; late stage enrichment highlighted hormone regulation, protein processing, nutrient response, intracellular homeostasis and HNF3B regulation—a well-established pathway essential for β-cell development^45,46^ etc, reflecting functional maturation. In contrast, differentiation toward SC EC cells showed early stage enrichment in vascular morphogenesis and GPCR signaling^47^, with the onset of external signal sensing, whereas late stage enrichment in hormone regulation, intracellular homeostasis, secretion, neuronal projection and sensory organ development^48^ likewise included HNF3B regulation indicated the maturation of secretory and sensory functions.

### Benchmarking interpolation and pseudotime inference of GeoBridge

We extensively benchmarked GeoBridge against several mainstream approaches, demonstrating its superior performance in terms of interpolation accuracy and pseudotime inference. First, we conducted held out interpolation experiments on the EMT^14^, EB^49^, and β cell^42^ datasets. Specifically, intermediate time points were masked, and GeoBridge was trained using the remaining data. The trained model was then used to reconstruct the masked samples. Interpolation accuracy was quantified using the Wasserstein distance between reconstructed and true distributions, with smaller values indicating higher accuracy. In PCA space, the reconstructed results showed substantial overlap with the real data (Fig. 4a–c), demonstrating that GeoBridge can accurately model continuous dynamics from discrete time point observations. Next, under the same held out setting, we compared GeoBridge with three representative neural ODE based methods—MioFlow^21^, TIGON^15^, and DeepRUOT^16^. After normalizing the accuracy results across experiments, we performed a t test to assess statistical differences. GeoBridge achieved significantly higher interpolation accuracy than competing methods (Fig. 4d; Supplementary Fig. S5a, Fig. S8). Because the optimal transport cost function in the flat forward manifold constructed by GeoBridge is Euclidean, the corresponding Kantorovich potential (KP) is theoretically linear with respect to real time parameter(Methods). Accordingly, the normalized KP values were defined as the inferred pseudotime. The accuracy of pseudotime inference was evaluated by the Pearson correlation between inferred pseudotime and true time normalized KP values. GeoBridge was then trained using these initialization labels, and after each training round the pseudotime was updated in the learned latent linear manifold. Taking the EMT dataset as an example, GeoBridge’s initial pseudotime already outperformed DPT^7^ and Palantir^8^ in correlation with true time, and iterative refinement further surpassed the well performing Slingshot^9^ method (Fig. 4e). We visualized comparisons between pseudotime inference results and the ground truth temporal order (Fig. 4f), along with the corresponding flattened manifold produced by GeoBridge (Supplementary Fig. S5b). Finally, we assessed whether GeoBridge trained purely on pseudotime labels could reconstruct biological dynamics comparable to those labels. For fair benchmarking against other pseudotime approaches, GeoBridge was trained without using real time labels. Specifically, initial pseudotime labels were computed on the original gene expression manifold using Euclidean distance based obtained when real time labels were used for training. Using the EMT dataset, we calculated gene wise correlations between dynamic expression profiles inferred under these two settings. The distribution across the top 1,999 variable genes showed that most genes clustered in high correlation regions (Fig. 4g), and typical EMT marker genes also exhibited highly consistent temporal trends (Fig. 4h). These results indicate that even for single cell RNA seq data lacking explicit temporal annotations, GeoBridge can reliably capture biologically meaningful dynamics, yielding reconstructions that closely match those derived from real time supervised models (Fig. S5 c–d).

**Figure 4.**
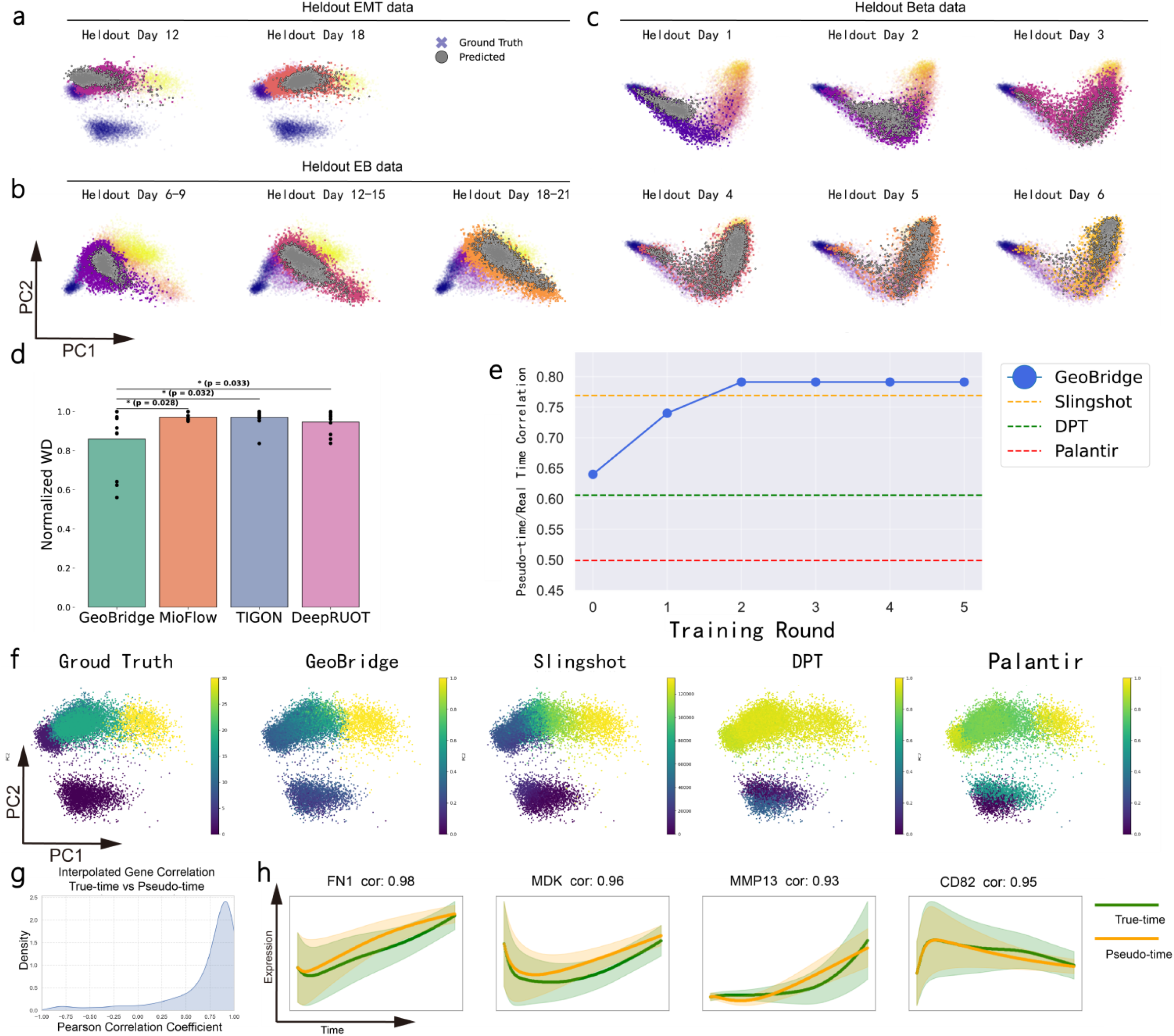
Benchmarking interpolation accuracy and pseudotime inference by GeoBridge. **a–c**, Held-out interpolation experiments on EMT, EB, and β-cell datasets, showing reconstructed intermediate time points (grey dot points) versus ground-truth (fork points) PCA projections. **d**, Comparison of normalized Wasserstein distances across methods (GeoBridge, MioFlow, TIGON, DeepRUOT) under held-out settings; GeoBridge achieves significantly higher interpolation accuracy (t-test). **e**, Pseudotime inference performance on the EMT dataset quantified by Pearson correlation between inferred pseudotime and true time labels; GeoBridge outperforms Slingshot, DPT, and Palantir, with iterative refinement further improving accuracy. **f**, Visualization of pseudotime inference results and ground-truth temporal order across methods. **g**, Distribution of gene-wise correlations between expression dynamics reconstructed using pseudotime-trained versus time-label-trained GeoBridge models, indicating strong consistency. **h**, Representative EMT marker gene trajectories (FN1, MDK, MMP13, SNAI2) inferred by the two training settings, showing highly similar temporal trends.

### Single-cell fate navigation in multi-target hematopoietic lineages

We applied GeoBridge to a human hematopoietic dataset^50^ comprising four time points and seven cell types, describing three major developmental lineage from hematopoietic stem cells (HSCs) toward neutrophil progenitors (NeuP), mast-cell progenitors (MasP) and megakaryocyte progenitors (MkP) (Fig. 5a). Unlike the independent training strategy used in previous results (Figs. 1–2), here we employed a unified, multi-target training scheme that jointly models all cell types using a single, more general INN model. This mapping to a flat forward manifold in which data evolve linearly over time yet are not constrained to a single direction of change. Within this forward manifold, we established optimal pairwise matching between day 2 and day 7 cells via Euclidean-cost optimal transport, and connected linear interpolation points by straight geodesics (Fig. 5b ②). The inverse mapping of these geodesics yielded nonlinear trajectories within the original gene-expression manifold (Fig. 5b ①), enabling the quantification of the gene dynamic driver index along the three distinct fate trajectories(Supplementary Fig. S6b-f). Intermediate interpolation results were visualized in PCA space to illustrate the complete temporal progression (Fig. 5c). In addition to generating navigation paths between any random pair of single cells(Supplementary Fig. S3h, S4h, S6g). We next performed a single-cell fate navigation (Path Compass) analysis in this multi-target system. For any given cell, GeoBridge enables generation of a geodesic path on the gene-expression manifold toward any chosen terminal state—potentially distinct from the cell’s predicted fate. Using Euclidean-cost optimal transport, we first calculated single-cell fate probabilities toward NeuP, MasP and MkP endpoints (Supplementary Fig. S6a, Methods). We then reconstructed mean geodesic trajectories from HSCs of NeuP fate to NeuP cells (green) and HSCs of MasP fate to MasP cells (red) (Fig. 5d). To test cross-fate navigation, HSCs originally biased toward the NeuP lineage were virtually guided toward MasP cells in the flat latent manifold by straight line geodesic, and the resulting navigation path (orange) was visualized in both the gene-expression and flat latent manifolds (Fig. 5d). To validate the biological plausibility of the navigated trajectory, we analyzed temporal dynamics of MasP marker genes (HDC, CPA3, MS4A2) and NeuP marker genes (ELANE, CSF3R, PRTN3) along both natural and navigated paths (Fig. 5e). The navigation path exhibited gene-expression patterns highly similar to the original MasP trajectory, indicating that the minimal-action path inferred by GeoBridge in forward-metric space represents a biologically feasible differentiation process. These results demonstrate that GeoBridge faithfully captures the geometric structure of the underlying hematopoietic fate manifold and supports single-cell-level navigation across multiple differentiation endpoints.

**Figure 5.**
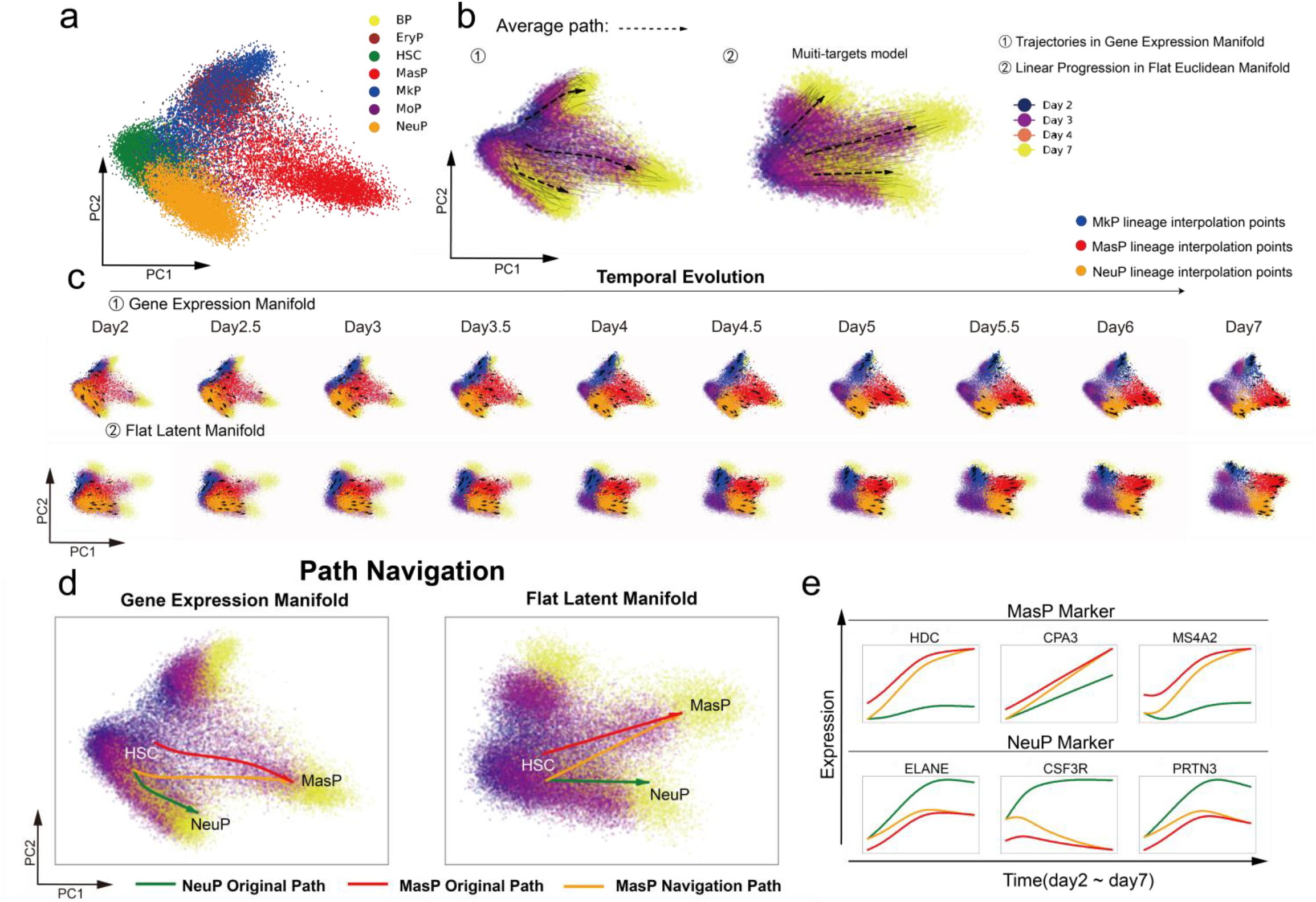
Single-cell fate navigation among multi-target hematopoietic lineages using GeoBridge. **a**, Human hematopoietic single-cell dataset spanning four time points and seven cell types, encompassing three developmental branches from hematopoietic stem cells (HSCs) to neutrophil (NeuP), mast-cell (MasP), and megakaryocyte progenitors (MkP). **b**, Illustration of the unified multi-target GeoBridge model: (①) 30 random selected nonlinear trajectories of each target in the original gene-expression manifold, and (②) corresponding linear geodesics in the flat forward (Euclidean) manifold connecting day 2 and day 7 cells. **c**, PCA-space visualizations of intermediate interpolation states between day2 and day7, showing continuous differentiation progression across MkP(blue), MasP(red) and NeuP(orange). **d**, Path Navigation for single-cell analysis. Mean geodesic trajectories are shown from HSCs to NeuP (green) and to MasP (red), with virtual cross-fate navigation directing NeuP-biased HSCs toward MasP cells (orange) plotted in both gene-expression and flat latent manifolds. **e**, Expression dynamics of representative lineage marker genes—MasP markers (HDC, CPA3, MS4A2) and NeuP markers (ELANE, CSF3R, PRTN3)—along original and navigated path. The navigated paths exhibit expression patterns consistent with the target lineage, demonstrating the biological plausibility and interpretability of GeoBridge-navigated trajectories.

## Discussion

GeoBridge provides a theoretically rigorous and geometrically elegant framework for reconstructing continuous cellular trajectories from discrete snapshots of single-cell transcriptomes based on our geodesic bridge theory (Theorem 2 in SI, or Equation 7), and further achieving single cell navigation between different cell states. Computationally by employing an invertible neural network to establish a bijective and isometric mapping or geodesic bridge/project between the original nonlinear manifold (gene-expression manifold) and an embedded linear manifold (flat Euclidean “forward” manifold), GeoBridge transforms nonlinear biological geodesics into tractable straight-line trajectories, enabling explicit modeling of dynamic optimal transport in high-dimensional space. This “geometric linearization” eliminates the need for computationally intractable geodesic estimation while ensuring a one-to-one correspondence between geodesics across manifolds, thereby characterizing the continuous evolution of cellular states as minimal-action trajectories. Importantly, during training GeoBridge establishes an optimal-transport mapping between the source and target domains, constructs linearly interpolated intermediate distributions, and minimizes the maximum mean discrepancy between these interpolants and the true data distributions. This design explicitly constrains the data distribution to evolve linearly and uniformly over time in latent forward manifold, functioning as an implicit noise-smoothing regularization that enforces temporal coherence without explicitly modeling stochasticity. Beyond reconstructing dynamic gene-expression trajectories, GeoBridge supports multiple downstream analyses, including pseudotime inference via the Kantorovich potential, single-cell fate probability estimation, dynamic driver-gene scoring, and in particular, trajectory navigation for virtual lineage switching. Furthermore, the geometric design of GeoBridge is inherently extensible: by replacing the INN with an invertible diffusion model^51^ and expanding the input from gene-expression vectors to images or spatial transcriptomic profiles, GeoBridge can be applied to the continuous reconstruction of discrete-time image sequences or 3D spatial data reconstitution^52–55^. This concept highlights the method’s broader potential for modeling spatiotemporal continuity and general dynamical systems beyond transcriptomics.

GeoBridge supports two complementary training paradigms: (1) supervised learning based on true temporal labels, and (2) unsupervised learning initialized with pseudotime estimates. The former provides explicit temporal anchoring and higher interpretability, whereas the latter leverages geometric self-consistency to achieve temporal alignment in unlabeled datasets, broadening the method’s applicability to usual single-cell datasets.

In addition, for multi-target developmental systems, we introduced both separate training and joint training strategies. Separate training more easily converges and accurately captures lineage-specific details, whereas the joint mode—by learning a shared latent linear manifold—produces more general representations transferable across lineages. These modes represent a trade-off between specificity and generality, and future hybrid strategies may yield both local accuracy and global geometric consistency.

GeoBridge can also extrapolate trajectories to predict future cellular states by extending straight-line paths in latent linear manifold. This approach accurately forecasts dynamics when evolution is approximately linear but performs sub-optimally for strongly curved or nonlinear trajectories (Supplementary Fig. S7). We attribute this limitation to insufficient information about future states. Biologically, cell fate is influenced not only by transcriptomic changes but also by epigenetic factors such as DNA methylation and chromatin accessibility (ATAC-seq). Incorporating multimodal datasets that integrate transcriptomic and epigenomic constraints could substantially enhance predictive accuracy and enable forward simulation of cellular fate decisions.

From a geometric viewpoint, GeoBridge learns the push-forward-metric distance ‖INN(x) − INN(y)‖^2^ between cells,. Because the INN mapping is constrained to be locally isometric, this learned metric equals the Riemannian distance on the original manifold and thereby encodes intrinsic biological regularities in state transitions. We postulate the existence of a universal biological distance function that quantitatively distance across any human cell types. Thus developing large-scale GeoBridge models trained on multi-tissue, multimodal datasets is thus not merely a technical pursuit but a necessary step toward uncovering geometric invariants that govern biological dynamics. Such foundation models may provide universal geometric representations enabling cross-dataset generalization, transferable inference, and interpretable modeling of any life’s continuous processes.

While GeoBridge captures intrinsic geometric regularities of state transitions, it currently neglects effects of growth, stochasticity, and cell–cell interactions. Future extensions incorporating mass-change, diffusion, and cellular interaction would further connect the learned geometry to realistic biological dynamics. In summary, GeoBridge reframes single-cell trajectory inference as a problem of intrinsic geometry, providing a principled and computationally efficient framework for reconstructing, interpreting, and navigating continuous cellular state transitions from sparse snapshot data, bridging the gap between discrete single-cell measurements and the continuous biological processes they represent.

## Methods

### Modeling Cellular Transitions via Dynamic Optimal Transport in GeoBridge

The goal of the dynamic optimal transport (OT) problem is to determine a flow field connecting a source probability distribution *µ* and a target distribution *v* such that the total action of the flow field is minimized:

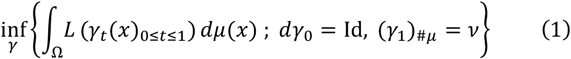

where *γ*_*t*_(*x*) denotes the trajectory starting from the initial position *x* over the time interval *t* ∈ [0,1].

The action functional of a trajectory, *L*(*γ*_*t*_(*x*)_0≤*t*≤1_), is defined in the Lagrangian form as:

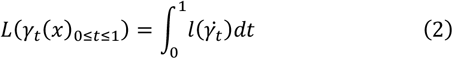

Where 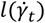 represents the instantaneous cost, *γ*_*t*_ is the position of the trajectory at time *t*, and 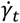 denotes its time derivative (i.e., velocity).

Suppose the mass is distributed on a Riemannian manifold, and let *γ*^*^ be the optimal trajectory that minimizes the above functional. If *γ*^*^ is a geodesic connecting *γ*_0_ and *γ*_1_ on the manifold, then a sufficient condition for the equivalence between the dynamic and static OT formulations is that the geodesic length equals the value of the distance function *c*(*x, y*) between the initial and terminal points:

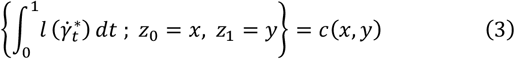

It follows that in the optimal transport problem, the choice of the cost function directly affects the structure of the optimal trajectory. In particular, when the cost function is the squared Euclidean distance, particles in the flow move at a constant speed along straight-line paths, the geodesic interpolation reduces to the McCann interpolation between the source distribution *µ* and the target distribution *v* and the geodesics degenerate to straight segments.

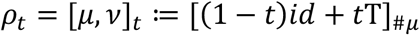

Based on the above formulation of the dynamic optimal transport problem, we next describe how this framework is adapted in our model to explicitly impose a constant-velocity straight-line constraint in the latent space. This constraint enforces that particles move along geodesics that degenerate into straight lines at a constant speed, ensuring a physically interpretable and regularized latent trajectory.

### Enforcing Predictable and Continuous Latent Dynamics

In this study, we mapped high-dimensional gene expression data to a latent space via an Invertible Neural Network (INN), to constrain time-series data to evolve along a straight-line trajectory at a constant velocity in the latent space, we introduce an interpolation constraint based on the Optimal Transport (OT) framework, and adopt the Maximum Mean Discrepancy (MMD) as the matching metric to form a differentiable training loss.

Specifically, for each sequence of input samples, we randomly select two time points as the starting and ending points of the trajectory. Between the corresponding latent vectors *z*_*s*_ and *z*_*t*_, we use the Euclidean distance as the cost function and compute a regularized transport plan matrix *P* via a differentiable OT solver denoted by get_ot_plan_func(·). The source data is then mapped to the target via the barycentric mapping:

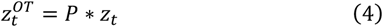

To enforce the constant-velocity linear constraint, we adopt a linear interpolation scheme between the starting latent vector *z*_*s*_ and its OT-mapped counterpart 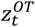, according to the relative time ratio:

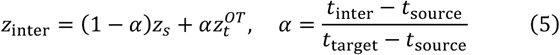

This ensures that the latent-space trajectory is linear with a constant speed (i.e., equal displacement over equal time intervals).

Interpolation is performed for each other time points, where *α* denotes the relative time proportion. The interpolated latent vectors are then passed through the inverse mapping of the INN to recover the corresponding data in the original space. Finally, the MMD between the generated samples and the true observations at the same time points is computed to measure the consistency between the generated linear trajectory and the actual trajectory, and the averaged MMD is used as the Constant-Velocity Linear loss:

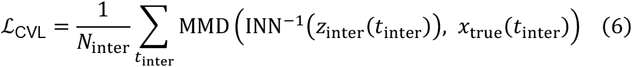

This approach explicitly constrains the dynamic trajectory in the latent space to follow a constant-velocity straight-line motion, effectively realizing a physically motivated trajectory assumption, while preserving full end-to-end differentiability for joint optimization.

### Establishing a Geometric Bridge Between Manifolds

We first obtain a geodesic in the latent space manifold as a constant-velocity straight segment. The latent features of intermediate points along this segment can be directly obtained via linear interpolation. These interpolated points are then mapped back to the original space through the inverse mapping of the INN to obtain their corresponding trajectories in the original manifold. However, such trajectories are not necessarily geodesics in the original manifold.

In the SI (Theorem 2), we show our geodesic bridge theorem, i.e. when the INN mapping of neighborhood between manifolds satisfies the local isometric constraint, there is a one-to-one correspondence between geodesics in manifolds *M* and *N*:

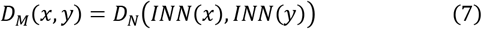

where *D*_*M*_ denotes the geodesic distance on the original manifold *M, D*_*N*_ denotes the geodesic distance on the mapped (latent) manifold *N* after the forward mapping, *x* and *y* are neighboring cells on the manifold, and *INN*(*x*) means to map the data *x* from *M* to *N*. INN is the diffeomorphism. Such a bijective project is the geodesic bridge, which maps geodesic of the two manifolds in a one-to-one manner. From an intuitive perspective, preserving the local distance relationships ensures that the local geometric structure is undistorted, which implies that the constant-velocity linear constraint flattens the manifold into the Euclidean space without introducing any local inversion or twisting.

### Preserving Intrinsic Biological Distances via Isometric Mapping

From Theorem 1 and Theorem 2, such a local isometry (Equation 11) also preserves geodesics: if *γ*(*t*) is a geodesic in 𝒩, its inverse image under *ϕ* is a geodesic in ℳ, which is called a bijective geodesic bridge or project in this work. This property is crucial for ensuring that trajectories in latent space correspond to geometrically consistent trajectories in the original space.

In practice, we aim to realize this condition in the INN framework by comparing nearest-neighbor distances in the original data space ℳ and in the mapped latent space 𝒩 = ϕ(ℳ). We adopt the standard local Euclidean approximation assumption: on a Riemannian manifold, for two sufficiently close points, the geodesic distance can be well-approximated by the Euclidean norm in local coordinates. Under this assumption,

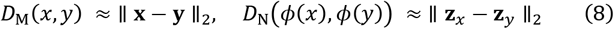

for local neighborhoods, where **z**_*x*_ = ϕ(**x**).

Let *m* and *n* index two distinct data samples, which we regard as two points on the original manifold ℳ (or its mapped counterpart 𝒩). The pairwise Euclidean distances are denoted as

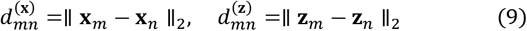

For each sample *m*, we compute the nearest-neighbor distance in ℳ as

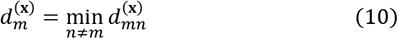

and in 𝒩 we take the distance to the corresponding nearest neighbor (found in ℳ):

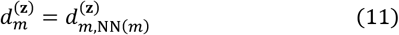

where NN(*m*) denotes the index of the nearest neighbor of **x**_*m*_ in ℳ.

We then form the ratio

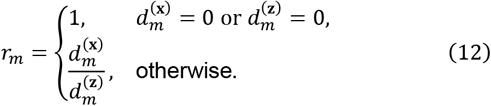

While the strict local isometry condition would require *r*_*m*_ ≈ 1 for all *m*, in training we relax this requirement to

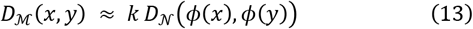

where *k* is an arbitrary global constant. This relaxation allows for a uniform scaling of geometry between the two manifolds. Under this relaxed constraint, what matters is that all *r*_*m*_ within a local neighborhood are close to the same constant *k*.

To enforce this, we define the isometry loss for a given time slice *t*_*i*_ as

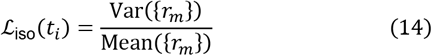

which is the coefficient of variation of the ratios {*r*_*m*_}. A smaller ℒ_iso_(*t*_*i*_) indicates that the ratios *r*_*m*_ are more concentrated around a common factor *k*, meaning the mapping preserves local distances up to an overall scaling. Averaging ℒ_iso_(*t*_*i*_) over all time slices yields the total local isometry regularization term used during training.

### Deep learning-based invertible solver with geometry-preserving regularization

Now we take everything together to describe our deep learning-based solver, where the mapping between the original manifold *M* and the latent manifold *N* is implemented by an Invertible Neural Network (INN). This framework is built on the Framework for Easily Invertible Architectures (FrEIA), which allows the rapid construction of complex invertible computation graphs from simple invertible building blocks. INNs guarantee exact forward and inverse computations by design, which is a exact bijectivity and diffeomorphism, preserving information in both mapping directions. The total loss function used to train the INN consists of Constant-Velocity Linear loss and Isometry loss:

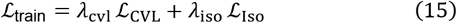

For model selection, we use a validation loss designed to measure the flatness of the latent manifold 𝒩. Given latent features *z* over time *t*, we compute the mean latent vector at each unique time, then calculate the mean-change vectors between all pairs of times. The validation loss is the mean pairwise squared distance between these mean-change vectors, with lower values indicating more constant latent-space velocities and hence a flatter latent geometry. This validation loss is computed every epoch and the model with the lowest validation loss is selected:

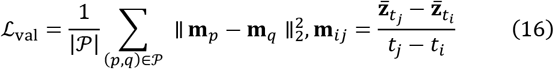

where 𝒫 is the set of all pairs of distinct elements in the collection *M*_diff_ = {**m**_*ij*_: *i* ≠ *j*}, and 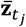denotes the mean latent vector at time *t*_*j*_.

### Heldout benchmarking setup

We benchmarked three representative dynamical optimal transport frameworks—MioFlow, TIGON, and DeepRUOT—on the EB, beta-cell differentiation dataset, following the evaluation protocol used for the EMT results. These three models respectively correspond to balanced OT, unbalanced OT, and regularized unbalanced OT formulations. All input gene expression data were preprocessed and embedded into a 50-dimensional PCA latent space, which serves as the common input representation for all baselines to ensure comparability.

For hyperparameters, we adopted the default configurations provided in the DeepRUOTv2 implementation, which already includes recommended settings for MioFlow, TIGON, and DeepRUOT. Specifically, we used the same learning rate, batch size, neural network backbone, and optimizer configuration across the three methods, except for the OT-related coefficients that are inherently required by each formulation (e.g., mass-variation penalties for unbalanced OT).

For evaluation, we computed weighted Wasserstein-1 (W1) distances for the two unbalanced-OT methods (TIGON and DeepRUOT) to appropriately account for variations in cell-state mass, whereas the standard W1 metric was used for MioFlow, which assumes a balanced-mass transport setting. This ensures that the evaluation metric is aligned with the underlying modeling assumptions of each OT formulation.

For GeoBridge, we performed held-out experiments on all three datasets using the same random seed (42) to ensure reproducibility. The INN module was initialized and parameterized with identical depth and width configurations which is same as used in the training stage. We also applied consistent hyperparameter settings across datasets, including learning rate, loss-function weighting, and batch size, which was scaled proportionally to the total number of samples in each dataset.

During evaluation, we adopted the standard W1 metric to quantify the discrepancy between the ground-truth cells and the GeoBridge-generated intermediate points, thereby measuring how closely the reconstructed trajectories align with the true cell distributions over time.

### Downstream analyses

### Reconstructing Continuous Developmental Trajectories in Data Manifold

In downstream analysis (Figure 1b(i, ii), to generate a continuous geodesic from discretely sampled time-series data, this study proposes a method based on optimal transport in the latent space combined with linear interpolation. Specifically, we first map the original high-dimensional data into latent vector representations using a pre-trained INN model. Then, source and target time points are selected from the discrete time point set, typically the minimum and maximum time points. Kernel density estimation (KDE) is used to construct the probability distributions of the starting and ending points in the latent space. Since the latent space is a flat space learned by the model, the cost matrix can be directly constructed using Euclidean distance. Similar to the training process, linear interpolation is applied between the source domain *z*_*s*_ and the barycentric mapping result 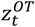 using the formula:

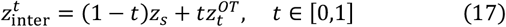

This generates latent vectors at intermediate time points, achieving uniform straight-line interpolation in the latent space. All interpolation results are then mapped back to the original data space via the inverse mapping of the INN, yielding reconstructed samples at continuous time points. The trajectory constructed in the original space lies on the geodesic of the original data manifold, corresponding to a straight-line geodesic in the latent space.

### Determination of Cell Fate

In this analysis (Figure 1b(iii)) we determined cells’ fate based on OT. Firstly, we construct a cost matrix between source and target time points in the latent space using the Euclidean distance, and solve for the OT plan *P*, where each row corresponds to a source cell and each column corresponds to a target cell. After normalizing each row of *P*, the entry *P*_*ij*_ represents the proportion of “mass” from the *i*-th source cell transported to the *j*-th target cell. For each predefined target cell type *c*_*k*_, let 𝒞_*k*_ be the set of column indices corresponding to that type. The total transport weight from the *i*-th source cell to type *c*_*k*_ is computed as:

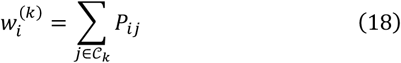

In the binary classification case (e.g., types *c*_1_ and *c*_2_), we define the fate tendency score as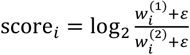, where *ε* is a small constant to avoid division by zero. The confidence of the fate assignment is defined as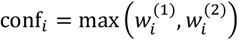

Given a tendency score threshold *τ*_score_ and a confidence threshold *τ*_conf_, we assign cell fates as follows: (1) if score_*i*_ > *τ*_score_ and conf_*i*_ > *τ*_conf_, the cell is assigned to type *c*_1_ ; (2) if score_*i*_ < −*τ*_score_ and conf_*i*_ > *τ*_conf_, it is assigned to type *c*_2_; (3) if |score_*i*_| ≤ *τ*_score_ with conf_*i*_ > *τ*_conf_, it is labeled as intermediate; and (4) if conf_*i*_ ≤ *τ*_conf_, it is labeled as low-confidence, indicating that its fate cannot be reliably determined.

In cases with more than two target cell types, each cell is assigned to the type *c*_*k*_ for which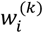 attains its maximum. This classification strategy jointly considers the relative tendency toward each fate and the absolute confidence of transport assignment, enabling robust and interpretable fate mapping at the single-cell level.

### Cell fate virtual navigation

To explore cell states that may exist but have not yet been experimentally observed, and to characterize the continuous dynamics of potential cross-lineage transdifferentiation processes, we developed a cell-fate virtual navigation module (Figure 1b(vi)) based on the GeoBridge framework. In brief, for any given pair of starting and target cells (or cell populations), we compute the geodesic path connecting them, which represents the theoretically minimal-energy and geometrically shortest transition trajectory between the two states.

To accurately capture the geometric structure of the cellular state space, all single-cell transcriptomic profiles are first mapped by the trained INN onto a forward Euclidean manifold. Each cell can be regarded as a point residing on a smooth manifold, where the Euclidean distance closely approximates the geodesic distance in the original gene-expression manifold. Within the GeoBridge formulation, a straight-line geodesic in the Euclidean manifold simultaneously corresponds to a minimal-action path; therefore, its length can be interpreted as the theoretically lowest-energy transition route.

For any specified pair of single cells, we can directly obtain the straight-line geodesic connecting them in the Euclidean manifold and then reconstruct their continuous trajectory on the original gene-expression manifold through the inverse mapping of the INN. For paired cell populations, KDE is used to estimate the probability density distributions of the source and target groups in the Euclidean space. We then compute a regularized optimal-transport plan via the Sinkhorn algorithm, obtaining a barycentric mapping that defines the straight-line geodesics connecting the two groups.

In practical applications, we first determine the fate label of each cell using the Cell-Fate Determination module of GeoBridge. Based on this, the virtual-navigation procedure computes the optimal transition trajectory from any given fate to any desired target fate and tracks the continuous evolution of gene expression along the inferred path. This process not only reveals potential differences in developmental potentials among distinct fates but also enables, in theory, the simulation of cross-lineage transdifferentiation, thereby providing a quantitative framework for identifying key regulatory genes and elucidating the mechanisms underlying cell-fate decisions.

### Inferring Developmental Time from Geodesic OT Potentials

In this study (Figure 1b(iv)), we propose a single-cell pseudotime construction method based on the Kantorovich potential from OT. Since the orignal data, after mapping through an INN, reside in a learned flat latent space satisfying Euclidean metric conditions, we adopt the squared Euclidean distance as the OT cost function. Under this assumption, the OT Monge map can be written as:

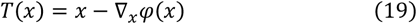

where *φ*(*x*) denotes the Kantorovich potential. The velocity *v* satisfies:

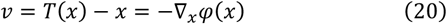

and assuming the velocity norm ∥ *v* ∥ is constant, the time derivative of −*φ*(*x*(*t*)) is given by:

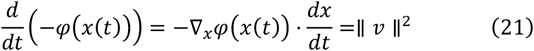

Integrating once with respect to *t* yields:

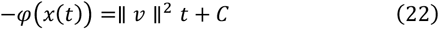

where the constant *C* is determined by the initial condition −*φ*(*x*(0)) at *t* = 0. This relationship demonstrates that −*φ*(*x*(*t*)) is strictly linearly proportional to real time *t*, with the proportionality coefficient equal to the squared velocity ∥ *v* ∥^2^ and *C* as the initial offset. For pseudotime analysis, the proportionality constant and offset do not affect the relative ordering of cells along the temporal axis; hence −*φ*(*x*) can be directly used as a quantitative pseudotime measure.

In practice, for unordered single-cell data, we first work in the original feature space and, based on known cell-type annotations or unsupervised clustering, we designate one cluster as the optimal transport target and treat all remaining cells as the source population. The regularized OT problem is then solved by the Fast Proximal Gradient Algorithm (FISTA) to obtain the initial −*φ*(*x*). This initial pseudotime is then used as a supervision signal to train the GeoBridge, mapping the data into the flat latent manifold. Finally, we rerun FISTA-OT in the latent space to update the Kantorovich potential and recompute −*φ*(*x*) as the final pseudotime, completing a training cycle. This method unifies the analytical equivalence between OT potential and temporal distance under Euclidean geometry with a learned space-flattening transformation, enabling stable and physically interpretable reconstruction of continuous single-cell trajectories.

### Identification of Dynamic Driver Genes

To identify potential driver genes along a specific cellular differentiation trajectory (Figure 1b(v)), we developed a method that combines trajectory interpolation based on OT with an INN. Following the interpolation procedure described in Equation (32), we first generate a continuous sequence of intermediate states in the latent space, starting from the initial state *z*_*s*_ and ending at the OT-derived mapped state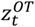.

To quantify the contribution of each gene to latent-space dynamics, we compute the mean velocity vector from the starting to the ending state:

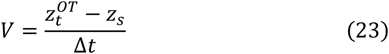

where Δ*t* denotes the temporal span of the trajectory. For an interpolated time point *τ*, let *X*(*τ*) in ℝ^*G*^ be the gene expression vector in the original feature space (*G* genes in total), and let *z*(*τ*) denote its corresponding latent-space representation. The weighted gradient of the latent features with respect to gene expression is then computed as:

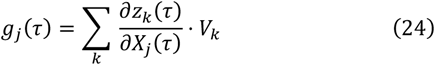

where *g*_*j*_(*τ*) quantifies the contribution of gene *j* at time *τ* to the trajectory’s velocity direction, *V*_*k*_ is the *k*-th component of the velocity vector in latent space, and the gradient term is obtained via the INN’s computational graph.

By aggregating these gradient-based contributions over all interpolated time points and averaging, we obtain the mean driver index for each gene:

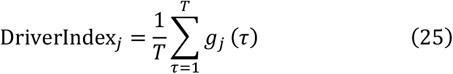

where *T* is the total number of interpolated time points.

Finally, all genes are ranked according to `DriverIndex`_*j*_, and the highest-scoring genes are selected as candidate driver genes most likely to exert pivotal regulatory influence along the given differentiation trajectory.

## Implementation Details

All models were implemented using PyTorch. The experiments were conducted on a workstation equipped with an NVIDIA GeForce RTX 3060 GPU. For a standard training configuration with a batch size of 3,000 cells, 2,000 highly variable genes (HVGs), and 500 epochs, the training process is typically completed in approximately 10 minutes.

## Data and code availability

All the datasets used and all source codes and models are publicly available at https://github.com/Zhu-JC/GeoBridge. All datasets used in this study were obtained from publicly available sources as described in the cited literature. The datasets had been preprocessed and standardized by the original authors, except for the HPSC dataset, which was further processed in this work using the Scanpy package. No additional preprocessing was performed in this work.

## Acknowledgements

This work was supported by National Natural Science Foundation of China (T2341007, T2350003, 12131020, 42450084, 42450135, 12326614, 12426310, T2542018 and 62002329); National Key R&D Program of China (2022YFA1004800, 2025YFF1207900, 2025YFC3409300); Science and Technology Commission of Shanghai Municipality (23JS1401300); Zhejiang Province Vanguard Goose-Leading Initiative (2025C01114); Shenzhen Medical Research Fund (E250200621); JST Moonshot R&D (No. JPMJMS2021).

## Notes

https://github.com/Zhu-JC/GeoBridge

### Competing Interest Statement

The authors have declared no competing interest.

