## Supplementary Information for "Generating and navigating single cell dynamics via a geodesic bridge between nonlinear transcriptional and linear latent manifolds"

#### Supplemental Note

#### Isometric geodesic theory

In GeoBridge, we first obtain a geodesic in the latent space manifold as a constant-velocity straight line via linear interpolation. These interpolated points are then mapped back to the original space through the inverse mapping of the GeoBridge to obtain their corresponding trajectories in the original manifold. However, such trajectories are not necessarily geodesics in the original manifold. We further proved that if the diffeomorphism(GeoBridge's INN model) between two manifolds is a local isometry, then there exists a one-to-one correspondence between their geodesic curves.

**- Lemma** for Components of the Pullback of a Metric Tensor: Let $M$ and $N$ be two smooth manifolds, with a smooth mapping $\phi: M \to N$. Suppose $N$ carries a metric tensor $g_{ab}$ with local components $g_{\mu\nu}$. We aim to compute the components of the pullback metric tensor $(\phi^* g)_{ab}$.

Consider a tangent vector $v^a \in TM$. Let $\{x^\mu\}$ and $\{y^\nu\}$ be local coordinate systems on $M$ and $N$, respectively. For the coordinate basis vector $\frac{\partial}{\partial x^\mu}$, and any scalar field $f$ on $N$, the definition of the pushforward gives:

$$\left(\phi_* \frac{\partial}{\partial x^\mu}\right)(f) = \frac{\partial}{\partial x^\mu}(\phi^* f) = \frac{\partial f(y)}{\partial x^\mu} = \frac{\partial y^\nu}{\partial x^\mu}\frac{\partial f}{\partial y^\nu} \tag{1}$$

Since $f$ is arbitrary, one has:

$$\left(\phi_* \frac{\partial}{\partial x^\mu}\right)^a = \frac{\partial y^\nu}{\partial x^\mu}\left(\frac{\partial}{\partial y^\nu}\right)^a \tag{2}$$

Now, for the pullback metric, by definition:

$$(\phi^* g_{ab}) v^a u^b = g_{ab}(\phi_* v)^a (\phi_* u)^b = g_{\mu\nu}\frac{\partial y^\mu}{\partial x^\rho} v^\rho \frac{\partial y^\nu}{\partial x^\sigma} u^\sigma \tag{3}$$

Since $v^a, u^b$ are arbitrary, it follows that:

$$(\phi^* g)_{ab} = g_{\mu\nu}\frac{\partial y^\mu}{\partial x^\rho}(\mathrm{d}x^\rho)_a \frac{\partial y^\nu}{\partial x^\sigma}(\mathrm{d}x^\sigma)_b \tag{4}$$

Then, we can prove our main theoretical results: Theorems 1 and 2 of bijective diffeomorphism and geodesic bridge theory, which ensure local isometric projects.

**-Theorem 1 :** Let $(M, g'_{ab})$ and $(N, g_{ab})$ be two Riemannian manifolds, and $\phi: M \to N$ a smooth mapping. If, for any two sufficiently close points $x, y \in M$, the geodesic distances satisfy

$$D_M(x, y) = D_N(\phi(x), \phi(y)) \tag{5}$$

then

$$g'_{ab} = (\phi^* g)_{ab} \tag{6}$$

**Proof**:

Choose local coordinates $\{x^\mu\}$ on $M$ and $\{y^\nu\}$ on $N$. Since condition 1 holds for any two sufficiently close points $p, q \in M$, take $p, q$ with small coordinate difference $v^\mu = x_2^\mu - x_1^\mu$.

Using the first-order expansion of the geodesic distance in $M$:

$$D_M(p,q) = g'_{\mu\nu} v^\mu v^\nu \tag{7}$$

For the image points under $\phi$, the coordinate difference is:

$$u^\mu = y_2^\mu - y_1^\mu = \frac{\partial y^\mu}{\partial x^\nu} v^\nu \tag{8}$$

Thus in $N$:

$$D_N\big(\phi(p), \phi(q)\big) = g_{\mu\nu} u^\mu u^\nu = g_{\mu\nu} \frac{\partial y^\mu}{\partial x^\rho} v^\rho \frac{\partial y^\nu}{\partial x^\sigma} v^\sigma \tag{9}$$

By the given condition $D^M(p,q) = D^N(\phi(p), \phi(q))$, we identify:

$$g'_{\rho\sigma} = g_{\mu\nu} \frac{\partial y^\mu}{\partial x^\rho} \frac{\partial y^\nu}{\partial x^\sigma} \tag{10}$$

which is precisely the pullback metric $(\phi^* g)_{\rho\sigma}$. ■

Then, we have our geodesic bridge theorem next, which is the theoretical basis of this work.

**-Theorem 2**: Let $(M, g'_{ab})$ and $(N, g_{ab})$ be Riemannian manifolds, with a smooth mapping $\phi: M \to N$. If $\gamma(t)$ is a geodesic in $N$ with respect to $g_{ab}$, then its pullback curve $\phi^{-1}(\gamma(t))$ (assuming it exists) is a geodesic in $M$ with respect to $g'_{ab} = (\phi^* g)_{ab}$.

**Proof**:

Let $\nabla_a$ be the torsion-free, metric-compatible connection on $N$ associated with $g_{ab}$. Define $\nabla'_a$ on $M$ via:

$$\nabla'_a T^{b_1 \cdots b_k}{}_{c_1 \cdots c_l} = \phi^*\big(\nabla_a (\phi_* T)\big)^{b_1 \cdots b_k}{}_{c_1 \cdots c_l} \tag{11}$$

First, we verify $\nabla'_a$ is torsion-free. For any vector fields $u'^a, v'^a$ on $M$, with pushforwards $u^a, v^a$ on $N$, the torsion tensor is:

$$T'^c{}_{ab} u'^a v'^b = u'^a \nabla'_a v'^c - v'^a \nabla'_a u'^c - [u', v']^c \tag{12}$$

By construction of $\nabla'_a$ and the compatibility of pushforward/pullback with tensor products and Lie brackets, we have:

$$T'^c{}_{ab} u'^a v'^b = \phi^*(u^a \nabla_a v^c - v^a \nabla_a u^c - [u, v]^c) = 0 \tag{13}$$

since $\nabla_a$ is torsion-free.

Second, $\nabla'_a$ is metric-compatible:

$$\nabla'_a g'_{bc} = \phi^*(\nabla_a g_{bc}) = 0 \tag{14}$$

Now, let $T^a$ be the tangent vector to a geodesic $\gamma(t)$ in $N$, satisfying $T^b \nabla_b T^a = 0$. Let $T'^a$ be the pullback of $T^a$ to $M$. Then:

$$T'^b \nabla'_b T'^a = \phi^*\big((\phi_* T'^b) \nabla_b (\phi_* T')^a\big) = \phi^*(T^b \nabla_b T^a) = 0 \tag{15}$$

showing that the pullback curve in $M$ is also a geodesic. ■

This theorem is our isometric geodesic theory, i.e. geodesic bridge theorem for GeoBridge method. During the training of the GeoBridge model, we introduced a local isometry regularization constraint to ensure that the reconstructed trajectory corresponds to the geodesic on the manifold.

#### Supplemental Information

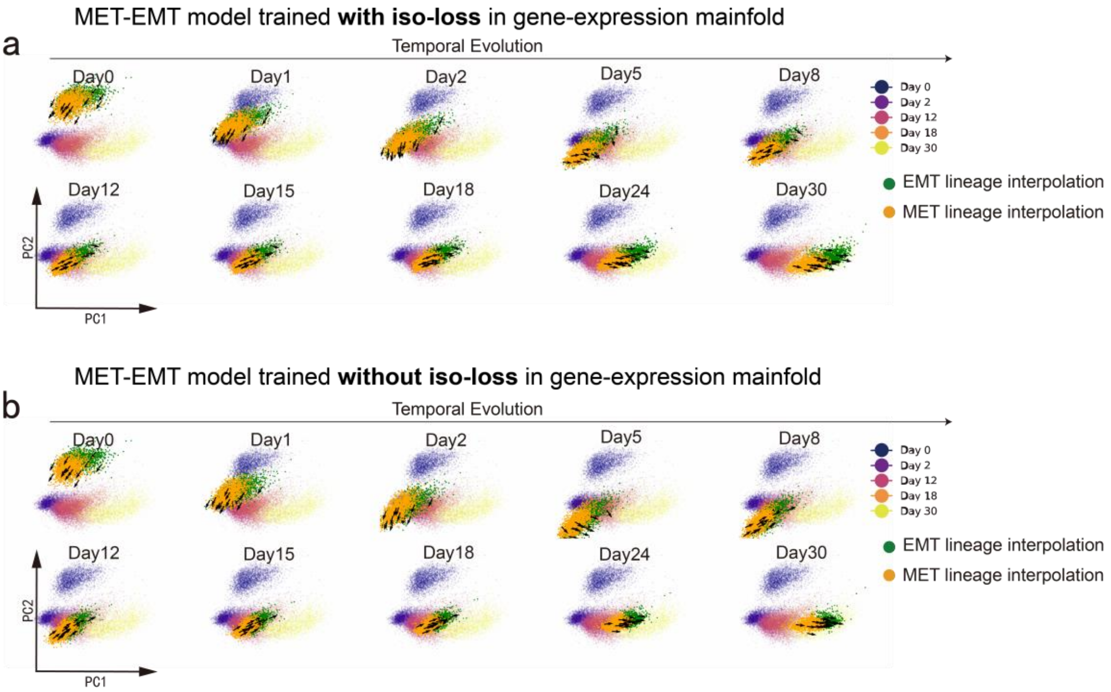


**Supplementary Figure S1 | Ablation study on the effect of the isometric loss. a**, Reconstruction of intermediate cellular states in the gene-expression manifold generated by a model trained with the isometric loss. **b**, Reconstruction of intermediate cellular states in the gene-expression manifold generated by a model trained without the isometric loss. The results show that models without the isometric loss produce interpolations that deviate substantially from the original data manifold, particularly at day 5 and day 8 as illustrated.

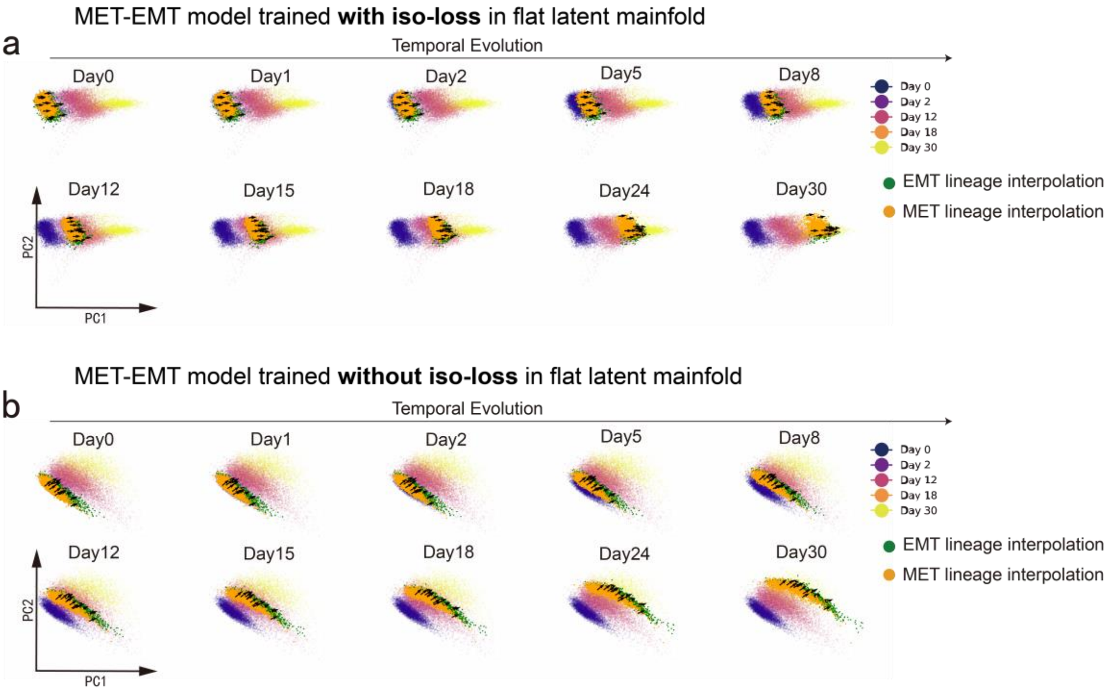


**Supplementary Figure S2 | Ablation study on the effect of the isometric loss in the flat latent manifold. a**, Reconstruction of intermediate cellular states in the flat latent manifold generated by a model trained with the isometric loss. **b**, Reconstruction of intermediate cellular states in the flat latent manifold generated by a model trained without the isometric loss. Both models produce interpolations that do not visibly deviate from the original data manifold in the flat latent space, indicating that a constant-velocity linear loss alone can yield a flat latent manifold geometry. However, without the isometric loss, straight geodesics in the latent space can be inversely mapped back to paths that are not geodesics on the original data manifold, potentially causing deviations from the true manifold in the original space.

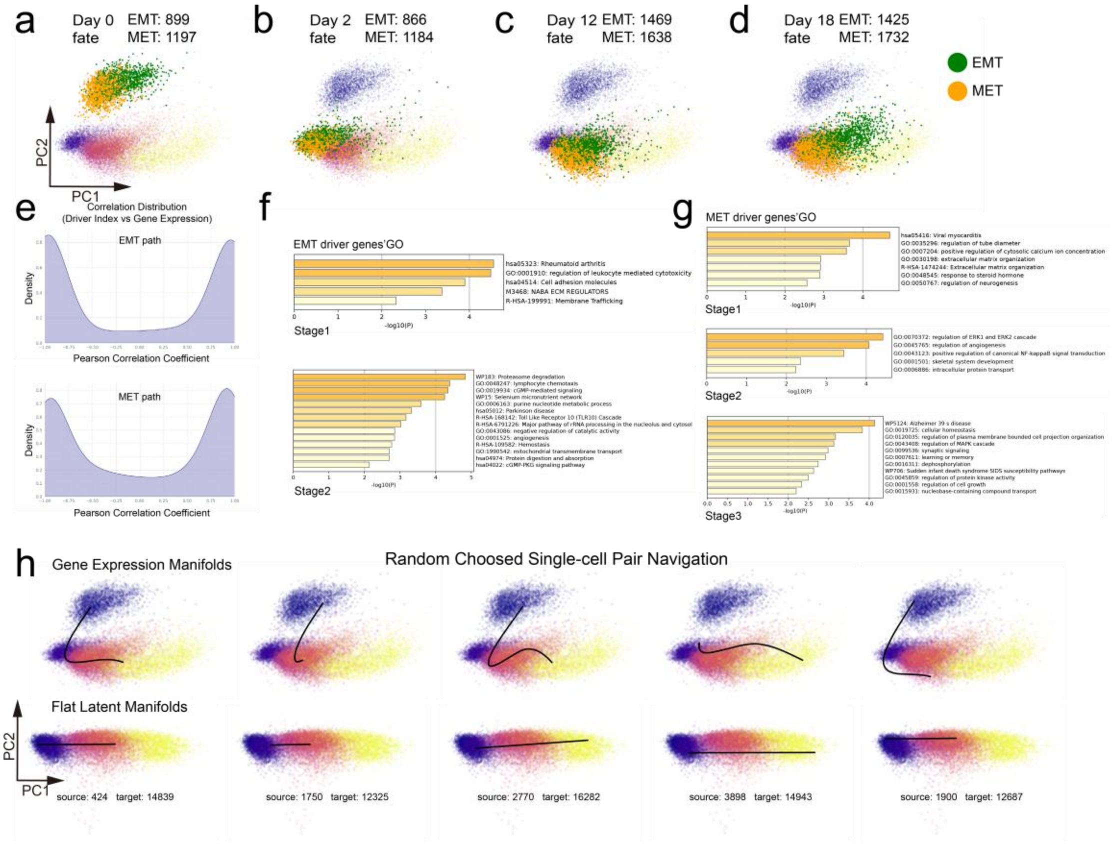


**Supplementary Figure S3 | Additional results of GeoBridge for EMT and MET progression datasets. a-d,** Cell fate determination results of individual cells in undifferentiated annotated cancer stem cells at day 0, day 2, day 12, and day 18. **e**, Pearson correlation coefficient between the gene dynamic driver index and gene dynamic expression over temporal evolution. **f-g**, EMT and MET Top 100 driver genes' GO analysis from metascape. **h**, Navigation trajectories for random choosed single-cell pairs, showing geodesic paths between two cells (top: gene expression manifold; bottom: flat latent space). Source and Target at the bottom indicate the indices of the two cells within the dataset.

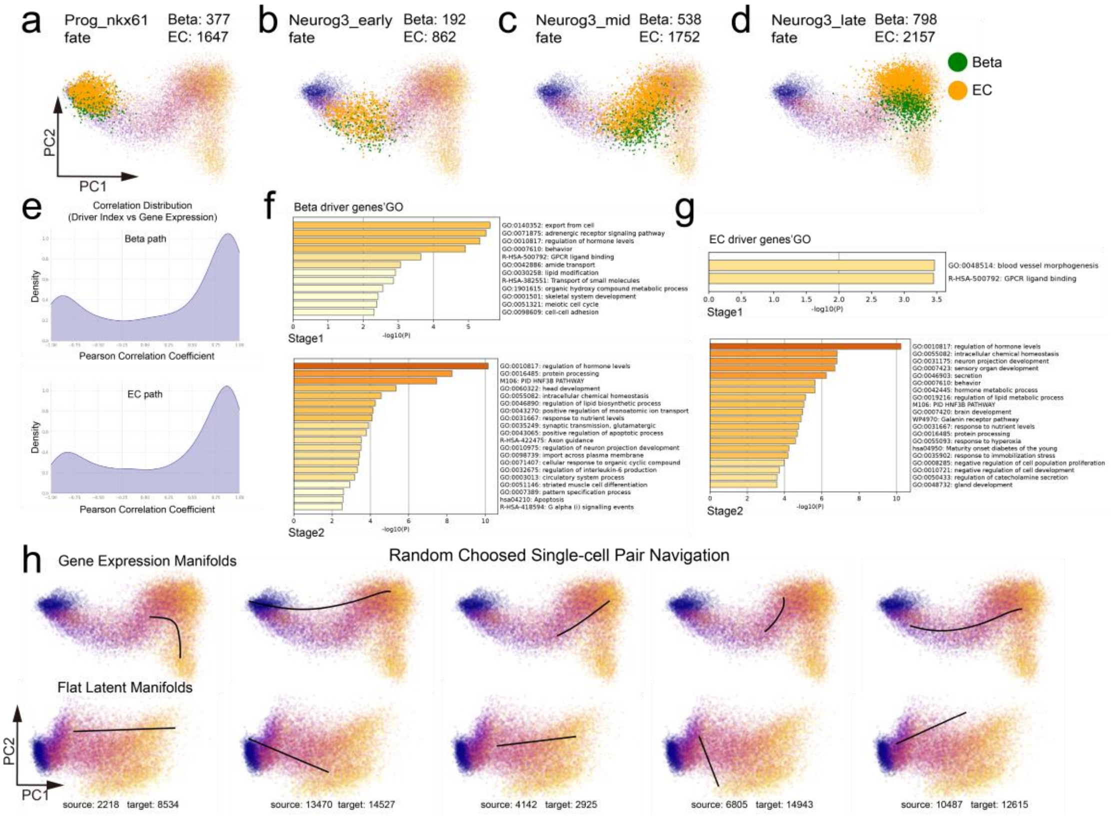


**Supplementary Figure S4 | Additional results of GeoBridge for pancreatic endocrine lineages datasets. a-d,** Cell fate determination results of individual cells in undifferentiated annotated HPSCs at Prog_NKX6.1, NEUROG3_early, NEUROG3_mid, and NEUROG3_late stage. **e**, Pearson correlation coefficient between the gene dynamic driver index and gene dynamic expression over temporal evolution. **f-g**, Beta and EC path Top 100 driver genes' GO analysis from metascape. **h**, Navigation trajectories for random choosed single-cell pairs, showing geodesic paths between two cells (top: gene expression manifold; bottom: flat latent space). Source and Target at the bottom indicate the indices of the two cells within the dataset.

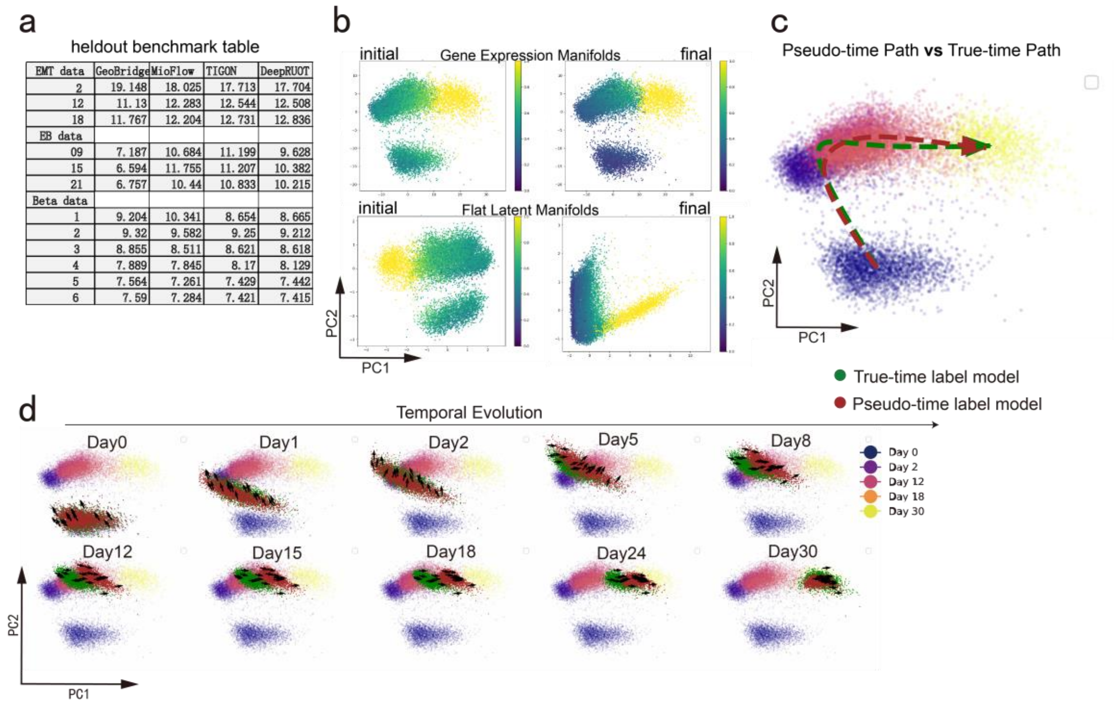


| EMT data | GeoBridge | MioFlow | TIGON | DeepRUOT |
|---|---|---|---|---|
| 2 | 19.148 | 18.025 | 17.713 | 17.704 |
| 12 | 11.13 | 12.283 | 12.544 | 12.508 |
| 18 | 11.767 | 12.204 | 12.731 | 12.836 |
| EB data | | | | |
| 09 | 7.187 | 10.684 | 11.199 | 9.628 |
| 15 | 6.594 | 11.755 | 11.207 | 10.382 |
| 21 | 6.757 | 10.44 | 10.833 | 10.215 |
| Beta data | | | | |
| 1 | 9.204 | 10.341 | 8.654 | 8.665 |
| 2 | 9.32 | 9.582 | 9.25 | 9.212 |
| 3 | 8.855 | 8.511 | 8.621 | 8.618 |
| 4 | 7.889 | 7.845 | 8.17 | 8.129 |
| 5 | 7.564 | 7.261 | 7.429 | 7.442 |
| 6 | 7.59 | 7.284 | 7.421 | 7.415 |

**Supplementary Figure S5 | Additional results of GeoBridge for heldout and pseudotime benchmarking. a,** Heldout results of four methods—GeoBridge, MioFlow, TIGON, and DeepRUOT—across three datasets. **b**, Initial and final pseudo-time inference results obtained through iterative updating in GeoBridge. **c**, Average trajectories constructed by GeoBridge using true-time label and iterative updated pseudotime label. **d**, Reconstruction of intermediate cellular states in the gene-expression manifold by true-time label model and pseudotime label model. The arrows indicate the velocities of ten randomly selected cells from each lineage. **c-d** Comparison of reconstructed continuous cellular dynamics, showing that a model trained with pseudotime labels achieves results highly similar to one trained with ground-truth experimental time labels.

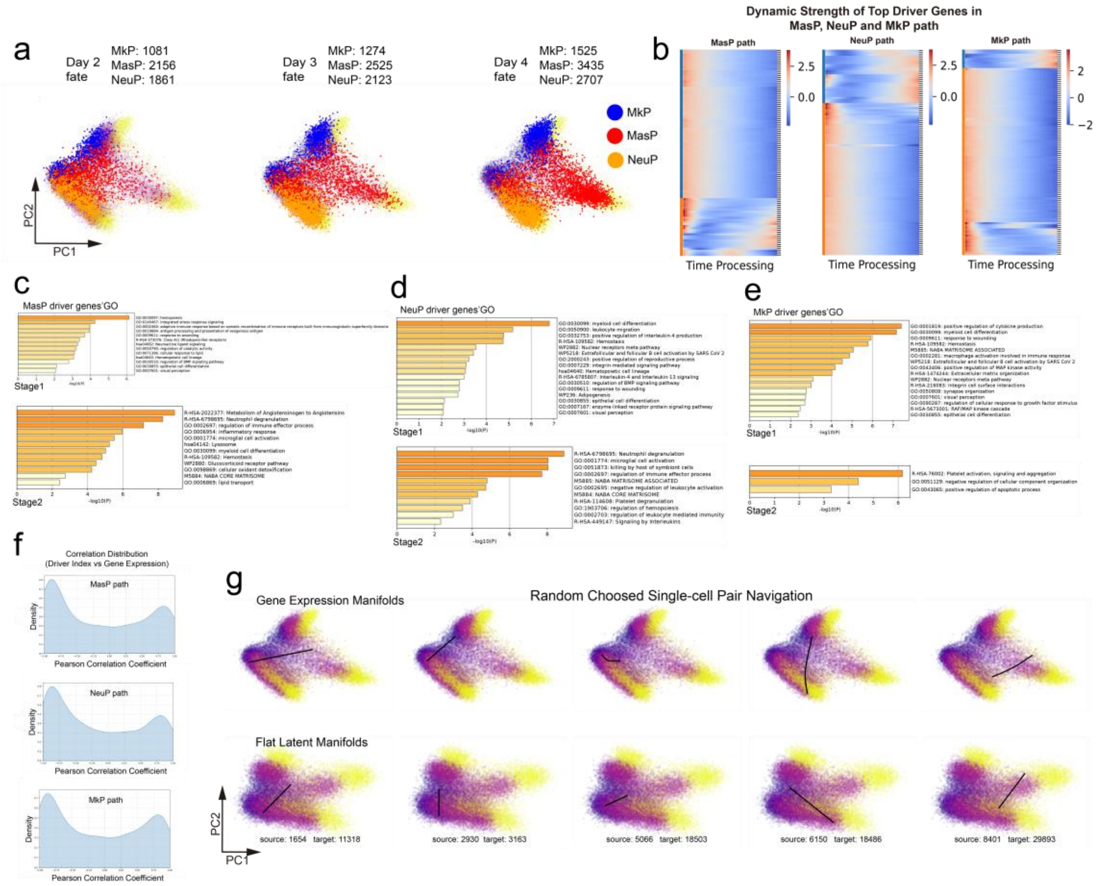


**Supplementary Figure S6 | Additional results of GeoBridge for multi target hematopoietic lineages datasets. a,** Cell fate determination results of individual cells in undifferentiated annotated HPSCs at day2, day3, and day4. b, Heatmap of Top 100 dynamic driver indices highlighting temporally distinct regulatory modules along MasP, NeuP, and MkP lineages. **c-e**, Beta and EC path Top 100 driver genes' GO analysis from metascape. **f**, Pearson correlation coefficient between the gene dynamic driver index and gene dynamic expression over temporal evolution. **g**, Navigation trajectories for random choosed single-cell pairs, showing geodesic paths between two cells (top: gene expression manifold; bottom: flat latent space). Source and Target at the bottom indicate the indices of the two cells within the dataset.

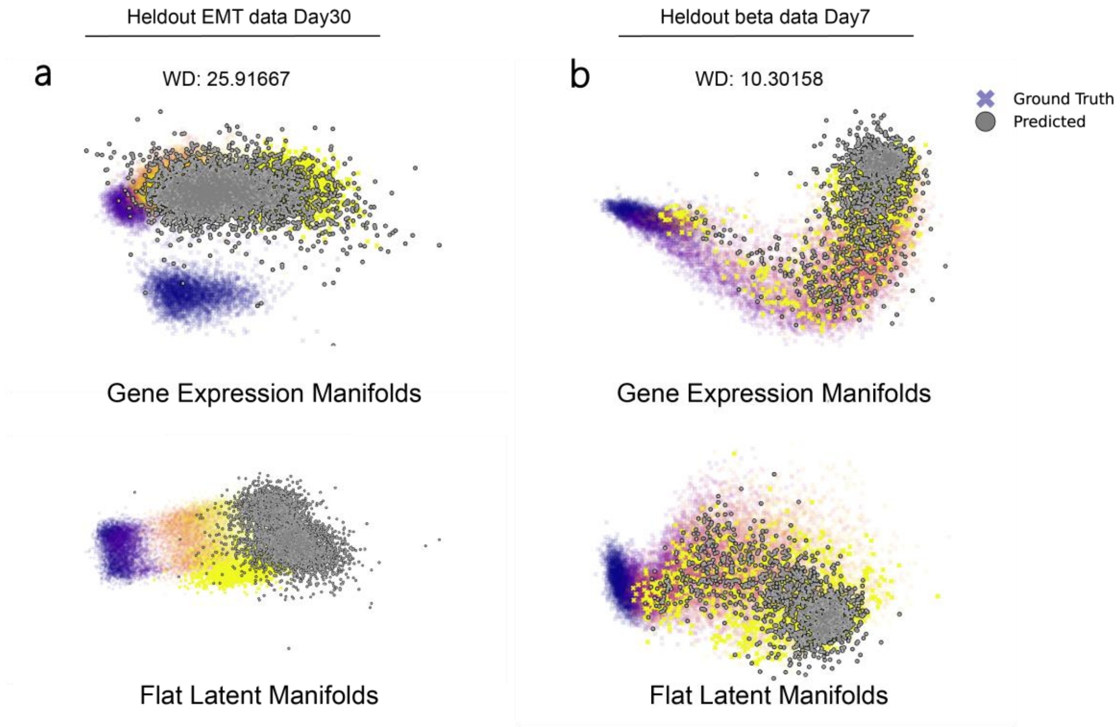


**Supplementary Figure S7 | Held-out prediction results for future time points. a**, Example from EMT dataset at day 30, where future state changes involve a pronounced deviation from prior trajectories, resulting in poorer prediction performance. **b**, Example from beta dataset at day 7, where future changes follow the existing trend with limited nonlinearity, leading to more accurate predictions.

### Heldout results in flat latent manifolds

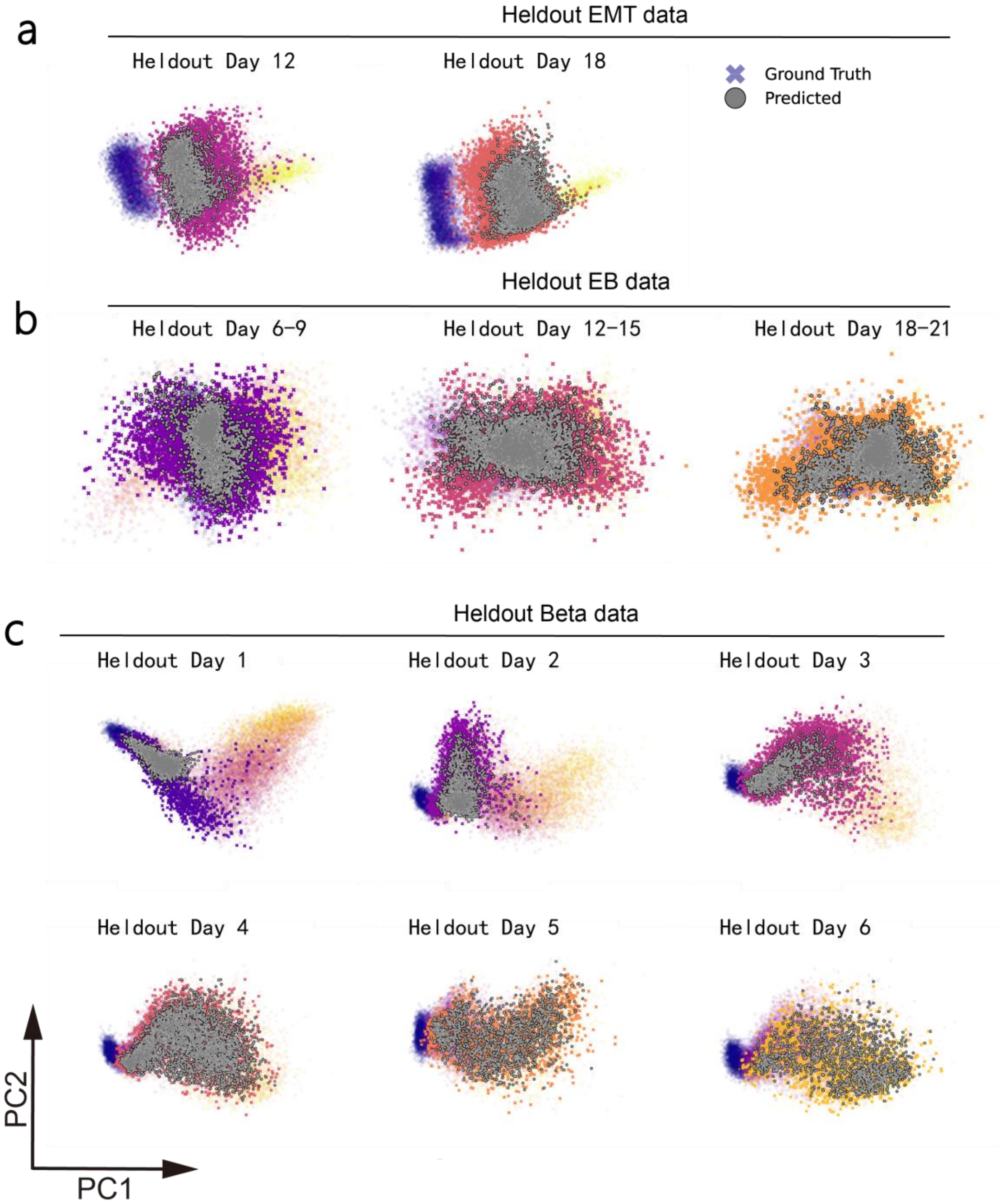


**Supplementary Figure S8 | Heldout interpolation results in flat latent results. a-c,** Heldout interpolation results visualized in the latent space of EMT, EB, and β-cell datasets. Generated points are highly overlapping with the ground truth, consistent with the corresponding visualizations on the gene expression manifold.

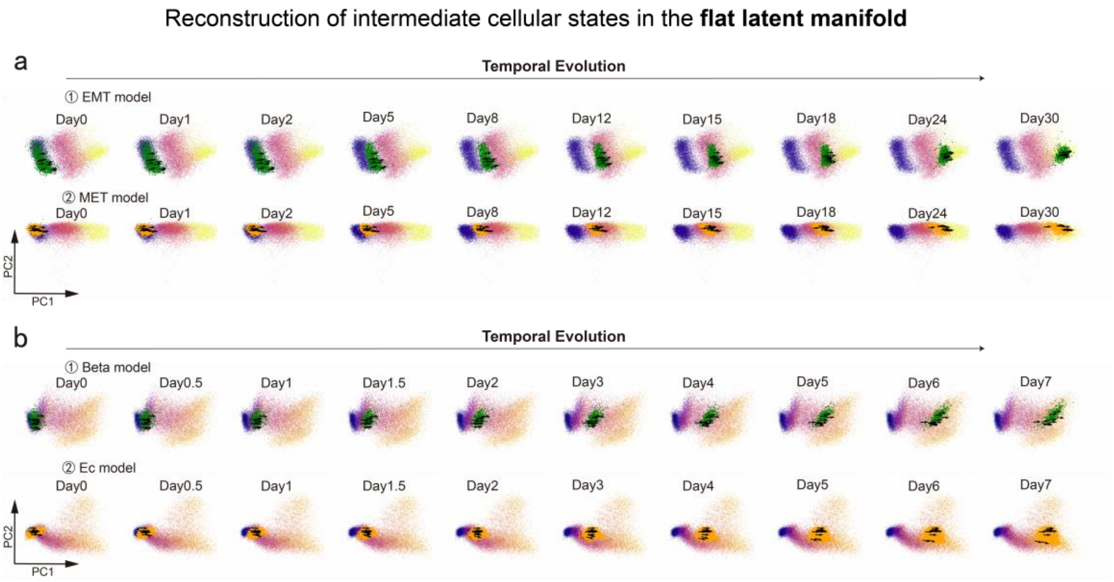


**Supplementary Figure S9 | Additional results of GeoBridge for reconstructing intermediate cellular states in the flat latent manifold. a,** Latent space dynamic interpolation along EMT and MET paths. **b,** Latent space dynamic interpolation along Beta and Ec paths. The arrows indicate the velocities of ten randomly selected cells from each lineage. The constant direction of the velocity vectors demonstrates uniform straight-line geodesics in the latent space.
